# Nanomechanics combined with HDX reveal allosteric drug binding sites of CFTR NBD1

**DOI:** 10.1101/2021.08.20.457065

**Authors:** Rita Padányi, Bianka Farkas, Hedvig Tordai, Bálint Kiss, Helmut Grubmüller, Naoto Soya, Gergely L. Lukács, Miklós Kellermayer, Tamás Hegedűs

**Affiliations:** Department of Biophysics and Radiation Biology, Semmelweis University, Budapest, Hungary; Faculty of Information Technology and Bionics, Pázmány Péter Catholic University, Budapest, Hungary; Theoretical and Computational Biophysics, Max Planck Institute for Biophysical Chemistry, Göttingen, Germany; Department of Physiology and Biochemistry, McGill University, Montréal, Quebec, Canada; ELKH-SE Molecular Biophysics Research Group, ELKH, Budapest, Hungary

**Keywords:** CFTR, F508 deletion, cystic fibrosis, atomic force spectroscopy, hydrogen-deuterium exchange, molecular dynamics simulations

## Abstract

Cystic fibrosis is most frequently caused by the deletion of F508 (ΔF508) in CFTR’s nucleotide binding domain 1 (NBD1), compromising CFTR folding, stability and domain assembly. The limitation of developing a successful therapy is due to the lack of molecules that synergistically facilitate folding by targeting distinct structural defects of ΔF508-CFTR. To improve drug efficacy by targeting the ΔF508-NBD1 folding and stability, and to study potential ΔF508-NBD1 allosteric corrector binding sites at the atomic level, we combined molecular dynamics (MD) simulations, atomic force spectroscopy (AFM) and hydrogen-deuterium exchange (HDX) experiments. These methods allowed us to describe unfolding intermediates and forces acting during NBD1 mechanical unfolding and to elucidate the stabilization mechanism of ΔF508-NBD1 by 5-bromoindole-3-acetic acid (BIA). An NBD1 region, including the α-subdomain, was identified as a potentially important participant of the first folding steps, characterized by non-native interactions of F508, thus destabilized in the deletion mutant. The instability was counteracted by the low-potency corrector BIA, increasing the mechanical resistance of the ΔF508-NBD1 α-subdomain, which was confirmed as a binding site by computational modeling and HDX experiments. Our results underline the complementarity of computational and experimental methods and provide a possible strategy to improve folding correctors.

## Introduction

Cystic fibrosis is a lethal disease caused by the functional defect of CFTR (Cystic Fibrosis Transmembrane Conductance Regulator) chloride channel in the apical membrane of epithelial cells [1, 2]. Biochemical, *in silico*, and structural studies of CFTR have contributed to understanding the effect of mutations ranging from misfolding to impaired regulation and channel gating [2–4]. CFTR is a member of the ABC (ATP Binding Cassette) protein superfamily that provides an ion conductance pathway through the cell membrane via two transmembrane domains (TMDs) each consisting of six TM helices [5–7]. CFTR possesses two nucleotide-binding domains (NBDs), which bind ATP, form a “dimer” via transient interactions, and regulate channel gating. An NBD is composed of a β-subdomain that binds ATP, and an α-subdomain that contains the ABC signature motif. The subdomains are not formed by sequential sequence regions but are intertwined (Fig. 1a). The binding and hydrolysis events are communicated towards the TMDs by the so-called coupling helices, which are the regions of intracellular “loops” that interact with the NBDs [5]. Most of these structural features have recently been confirmed by cryo-electron microscopy [6–9]. Mutations cover every region of the protein, and many of them are located in the N-terminal nucleotide binding domain, NBD1 [4].

**Fig. 1:**
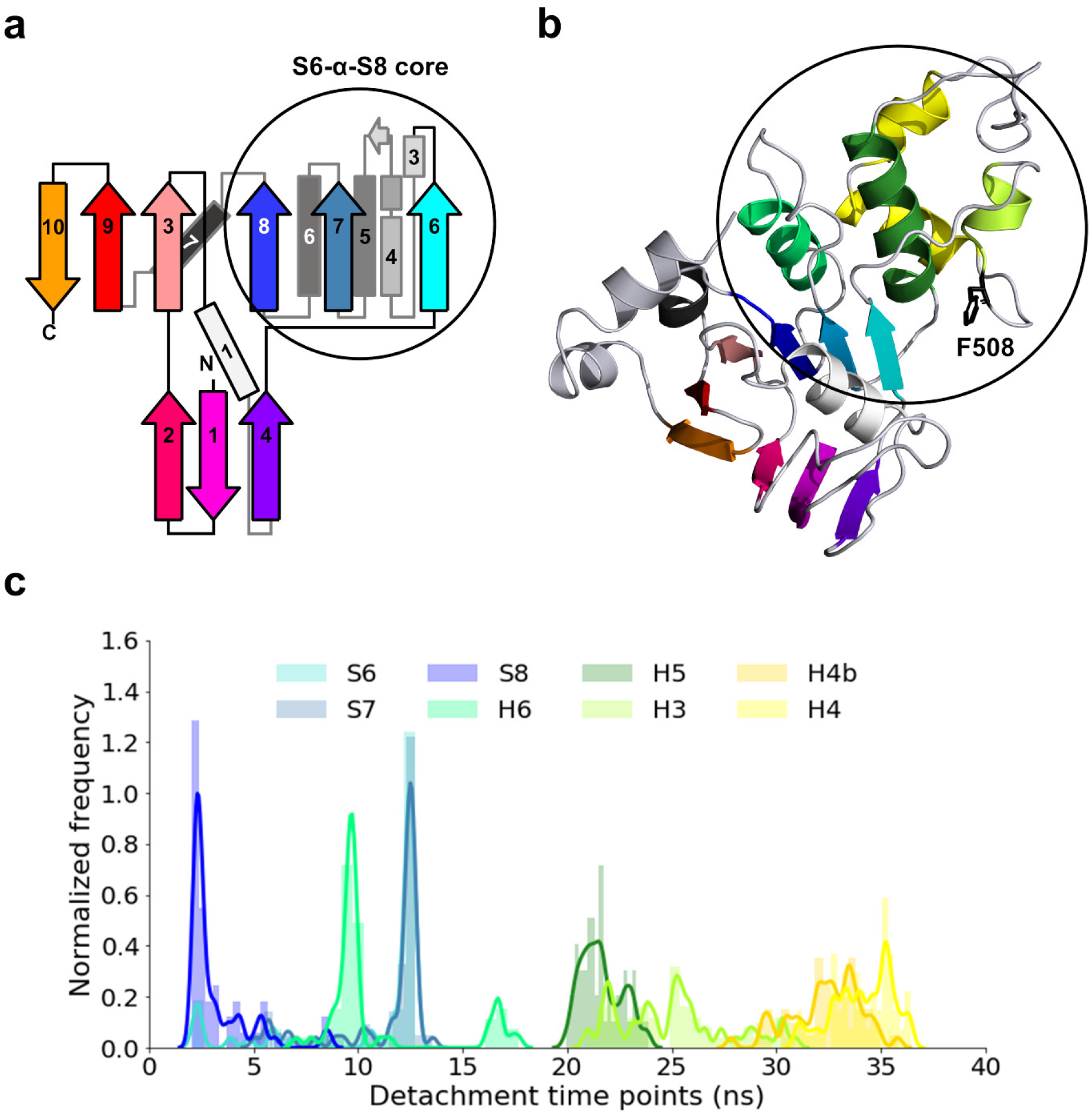
Organization of CFTR NBD1 structure and secondary structural element detachment. Topology (**a**) and structure (**b**) of CFTR NBD1. β-strands are colored, and α-helices are marked in shades of grey. Black circles label the S6-α-S8 core region. Green and yellow helices: α-subdomain. (**c**) Distribution of the detachment time points of each secondary structural unit calculated from all MD trajectories with the wild type S6-α-S8 core (pulling velocity was 1 m/s).

Sequence alterations in CFTR likely affect protein function, folding or stability. The effect of a mutation, such as the most frequent deletion of F508 residue (ΔF508 or F508del) located in NBD1, is usually complex and alters many of the above processes [4]. This deletion impairs the global domain-domain assembly of CFTR and affects the interactions between transmembrane domains, NBD1/TMDs, and NBD1/NBD2 [10–12]. A couple of specific amino acid interactions have been revealed to play an important role in CFTR domain assembly.

We have shown earlier that the hydrophobic side chain of F508 is crucial for the NBD1/CL4 interface, and the hydrophobic pocket of NBD1 around this amino acid could be a primary drug target for corrector development [5]. Furthermore, the NBD1/TMD assembly problem affecting global CFTR folding originates from local effects of ΔF508 in NBD1, and in the presence of this deletion NBD1 has shown decreased thermal stability [13–15].

Detailed studies of isolated wild type and ΔF508-NBD1 demonstrated a substantially decreased melting temperature of the mutant domain [13, 16] while suggesting a similar folding pathway compared to the wild type [17]. It was also proposed that a partially unfolded state is responsible for the aggregation propensity of NBD1. As primary objective in drug development the restoration of the ΔF508-CFTR folding was advocated by either exclusive correction of the NBD1 stability [18] or targeting both the NBD1 and NBD1-CL4 interface instability [19, 20]. Both methods are likely effective only on an already folded NBD1 subpopulation. Notably, the only approved drug that can target the isolated NBD1 was suggested to stabilize an (un)folding intermediate and not the native fold [21].

Early in CFTR research, altered folding of ΔF508-NBD1 has been demonstrated, and it was shown that the deletion affects a folding step prior to the ATP binding site formation [22, 23]. Studies on the co-translational NBD1 folding revealed that the synonymous codon for I507 upon F508 deletion results in lowering the speed of translation and has an effect on CFTR conformation [24, 25]. The folding of the nascent NBD1 polypeptide chain emerging from the ribosome was also extensively investigated using FRET [17, 26, 27]. In contrast to the previous results, these studies indicated that NBD1 folding starts on the ribosome and suggested that the ΔF508 defect occurs late in the NBD1 folding pathway.

An effective correction of any CF mutation must correct all the impaired steps of CFTR folding and/or its function. The first success in CF drug development is represented by Ivacaftor (Kalydeco, VX-770) which targets the natively folded G551D CFTR gating mutant [28]. The cryo-EM structure of the drug-bound WT CFTR complex provided the first hint for Ivacaftor’s mechanism of action at atomic resolution [29], launching the design of novel scaffolds for more efficient drugs. This would not only be important for G551D targeting, but also for other mutants with impaired function since the efficacy of Ivacaftor depends on the specific mutation type in the patient (https://pi.vrtx.com/files/uspi_ivacaftor.pdf). Lumacaftor (VX-809) has also emerged as a potential corrector of the ΔF508-CFTR folding defect [30, 31]. Although VX-809 has a low efficacy in restoring the ΔF508-CFTR folding [19], serious efforts have been devoted to identify its binding site [32]. Experiments suggested that VX-809 exerts its action on the TMD1 and TMD1/NBD1 interaction and also binds to the cleft formed by the C-terminal helices and β-strands S3, S8, and S9 [32–34].

The recently approved treatment for F508del, Trikafta, is a combination of VX-661 (Tezacaftor), VX-445 (Elexacaftor), and the gating potentiator VX-770 (Ivacaftor) [31]. VX-661 is a Type I corrector that restores the NBD1/TMDs interface. VX-445 was shown to be a Type III corrector that acts on NBD1 [21, 35]. Long-term studies of Ivacaftor indicated that adult G551D patients experience bacterial infections and progressive loss of lung function in spite of an initial partial normalization of the lung function [36, 37]. As the Trikafta treatment also exhibits only partial restoration of lung function, a long-term decline in effectivity of this drug combination can also be expected [31].

Since the lack of a type III corrector, which promotes NBD1 folding, was likely the bottleneck in developing an effective combination therapy such as Trikafta [31], improving existing and developing novel, more efficient folding correctors would be critical for long-term CF therapy. Therefore, the major objective of our study was to gain insights on NBD1 folding and its folding intermediates at a higher resolution compared to earlier studies. To identify unfolding pathways and intermediates, we performed both force-probe molecular dynamics simulations and force spectroscopy experiments on the wild type and ΔF508-NBD1. Since BIA (5-bromoindole-3-acetic acid) has been suggested as a type III corrector, we also investigated its effect on unfolding and determined its binding site using hydrogen-deuterium exchange (HDX).

## Results

### Differences in unfolding MD simulation with ΔF508 and WT S6-α-S8 cores

The structure of the CFTR NBD1 domain was investigated using a combination of experimental and simulation methods. To help the interpretation of the experimental AFM results, first we obtained a set of possible unfolding pathways that can be correlated with the experiments. Since pulling a large system such as the 250 a.a. long CFTR NBD1 in regular steered molecular dynamics simulations (MD) is highly limited, an unfolding pathway set of the whole WT CFTR NBD1 was collected using an all-atom Gō model (SMOG [38]). To identify the unfolding steps of NBD1, we analyzed the hierarchy in the detachment of secondary structural elements (SSE), which was calculated by the fraction of native contacts (Q) as a function of pulling time (Supplementary Fig. 1-2). We found that the unfolding of NBD1 consisted of two parts. The unfolding always began with the unfolding of the β-sheet subdomain followed by the α-helical subdomain.

Since experiments have indicated the S6-α-S8 core (the center of the α-helical subdomain, a.a. 487-603, Fig. 1) to be crucial for NBD1 folding [17, 39] we studied the unfolding of the S6-α-S8 core region, which occurs in the late steps of the unfolding pathway, in fully solvated atomistic force field pulling MD simulations for high resolution and accuracy. In the MD simulations of the WT and ΔF508 S6-α-S8 core we analyzed the order and the timing of SSE detachments and the rupture forces at which these detachments occur. The order of SSE detachments was determined by monitoring the fraction of the native contacts as was done for the Gō simulations (Supplementary Fig. 1). Detachment sequences of the secondary structure units were determined in each simulation, and the frequencies of the resolved pathways were calculated (Fig. 2a-c). In order to simplify the comparison between the WT and mutant pathways, we separated the S6-α-S8 core unfolding into two stages. In the first stage (Fig. 2b), the β-strands S6, S7 and S8 and the helix H6 unfold. Most frequently S8 decoupled in the first step, followed by H6 unfolding and the concurrent detachment of S6 and S7. The frequency of this pathway is much lower in ΔF508 S6-α-S8 core than in WT (54%, 27 out of 50 versus 78%, 39 out of 50; p < 0.05, χ^2^ test). The main divergence between the WT and ΔF508 protein was the different timing of the H6 detachment. An increased frequency of the pathways, in which the detachment of H6 occurs after the decoupling of S6 and S7, can be observed in ΔF508 S6-α-S8 core (ΔF508: 40%, WT: 22%; p=0.0517, χ^2^ test). This could be caused either by weaker binding of S6 or by stronger binding of H6 to the folded part of the core when compared to the WT. Comparing the S6 rupture forces of WT and mutant shows that in the case of ΔF508 a greater proportion of S6 detached at lower forces, while the distribution of H6 rupture forces were unchanged (Fig. 2d), indicating that the mutation weakened the S6 interactions. At the second stage of the S6-α-S8 core unfolding, the α-helices H5, H4, H4b and H3 detached (Fig. 2c). H5 and H3 in the WT core showed a synchronized unfolding slightly more frequently than in the ΔF508 core (20%, 10 out of 50 versus 6%, 3 out of 50; p=0.074, χ^2^ test). Comparing the time points of the SSE detachments of the WT and the mutant core, we observed faster detachment of secondary structural units (S6, H4 and H5) in the mutant compared to WT (Supplementary Fig. 3-4).

**Fig. 2:**
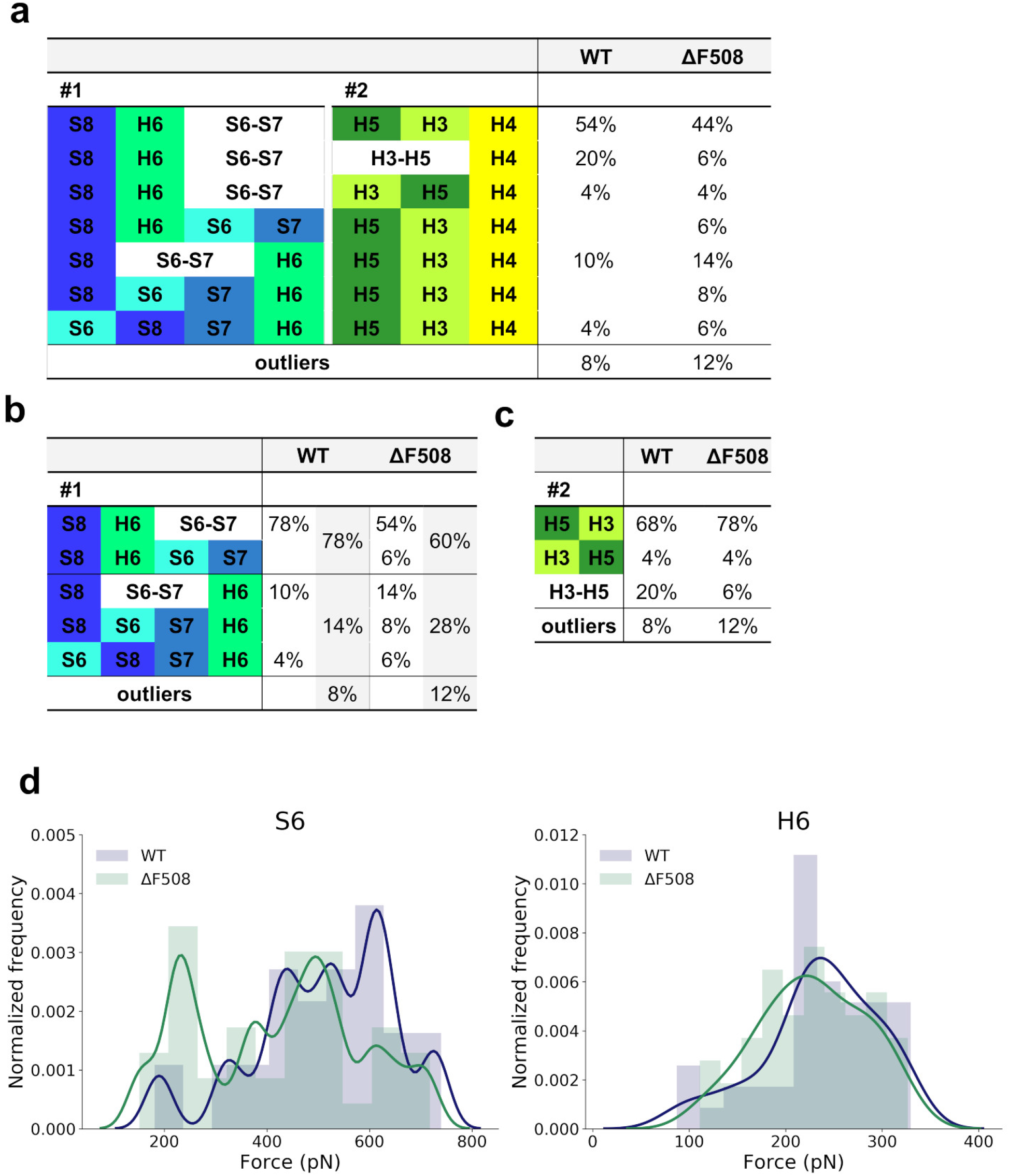
Alternative unfolding pathways of S6-α-S8 core in pulling simulations. (**a**) Pathways were determined by the detachment sequence of secondary structure units. Synchronized unfolding of two elements is marked by hyphenation and enclosing them in one cell. H4 labels both the H4 and H4b helices since they always unfold at the same time. (**b**) Summary of the pathway frequencies of the first stage of the S6-α-S8 core unfolding. Helix H6 unfolds last in all outlier pathways. (**c**) Summary of the pathway frequencies of the second stage of the S6-α-S8 core. (**d**) Unfolding force distribution of secondary structural elements S6 and H6.

To assess the effect of speed on the unfolding forces, we performed pulling simulations also at a lower speed, which resulted in smaller forces and loading rates (Supplementary Fig. 5-6). Nevertheless, the main pathways remained the same with somewhat shifted ratios; and some SSE detached earlier in the mutant than in the wild type (Supplementary Fig. 4 and 7).

### An increased number of non-native contacts are characteristic for late WT NBD1 intermediates

To characterize the details of different aspects of the unfolding steps, intermediate structures of the S6-α-S8 core from our MD simulations were determined and analyzed by using a special contact-based metric [40], which is efficient to compare highly different structures developing along pulling trajectories. Accordingly, the conformations from each pulling simulation were clustered separately based on contact RMSD. Then, the centroids of these clusters were pooled as intermediates, which were clustered again to yield all observable intermediates from every simulation. We identified four intermediate clusters describing the structural changes during unfolding in the case of both constructs (Fig. 3). However, there is an intermediate structure forming a well-defined wild type cluster (WT cluster #3), which was not observed for the ΔF508 S6-α-S8 core. Instead, the mutant conformations in the corresponding period (18-25 ns) changed their structure continuously into the totally unfolded state and were clustered into the last unfolding group.

**Fig. 3:**
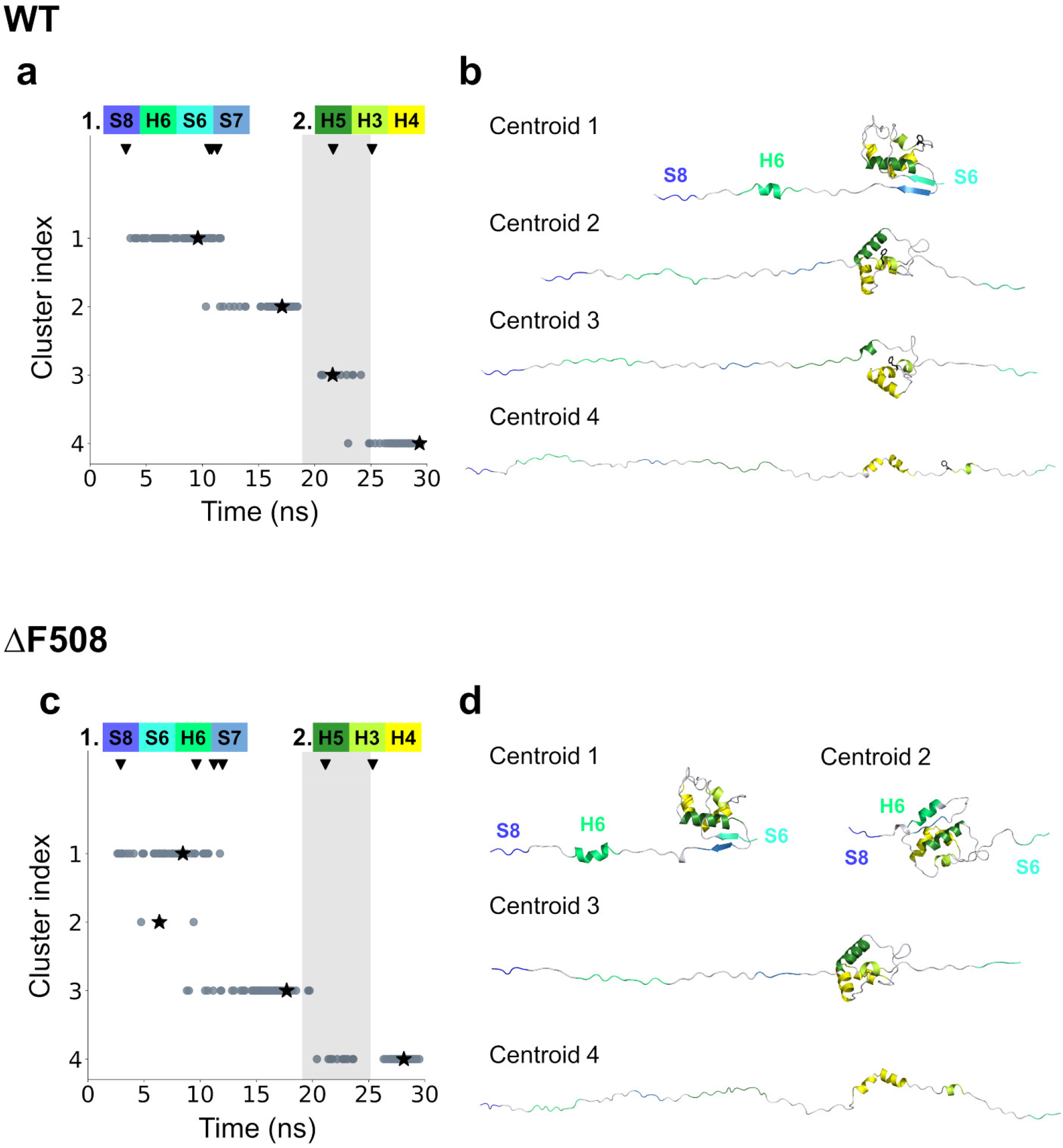
Clusters of S6-α-S8 core intermediates during unfolding. Intermediate structures from WT (**a**) and F508del (**c**) pulling simulations were clustered using contact RMSD as a pairwise distance metric. Cluster centroids are indicated by stars and their structures are shown on the right (**b, d**). The grey area highlights the cluster with intermediate structures present in the wild type core but not in the mutant core. Arrowheads mark the averaged detachment time points of each secondary structural element calculated based on its fraction of native contacts.

To analyze the intramolecular interactions contributing to the formation of the well-defined intermediate structure observed in the WT cluster #3, the native and non-native contacts were calculated during unfolding. As expected, the number of native contacts decreased monotonically during unfolding (Supplementary Fig. 8c) and appeared very similar for both constructs. In contrast, the number of non-native contacts increased at the beginning of pulling simulations (Fig. 4a and Supplementary Fig. 8e), probably reflecting the equilibration of the structure under pulling conditions, then exhibited a decreasing trend. Interestingly, a secondary increase was observed in conformations between 18 and 25 ns and it was more pronounced in the WT than in the mutant, coinciding with an unfolding intermediate state detected only in the WT (WT cluster #3).

**Fig. 4:**
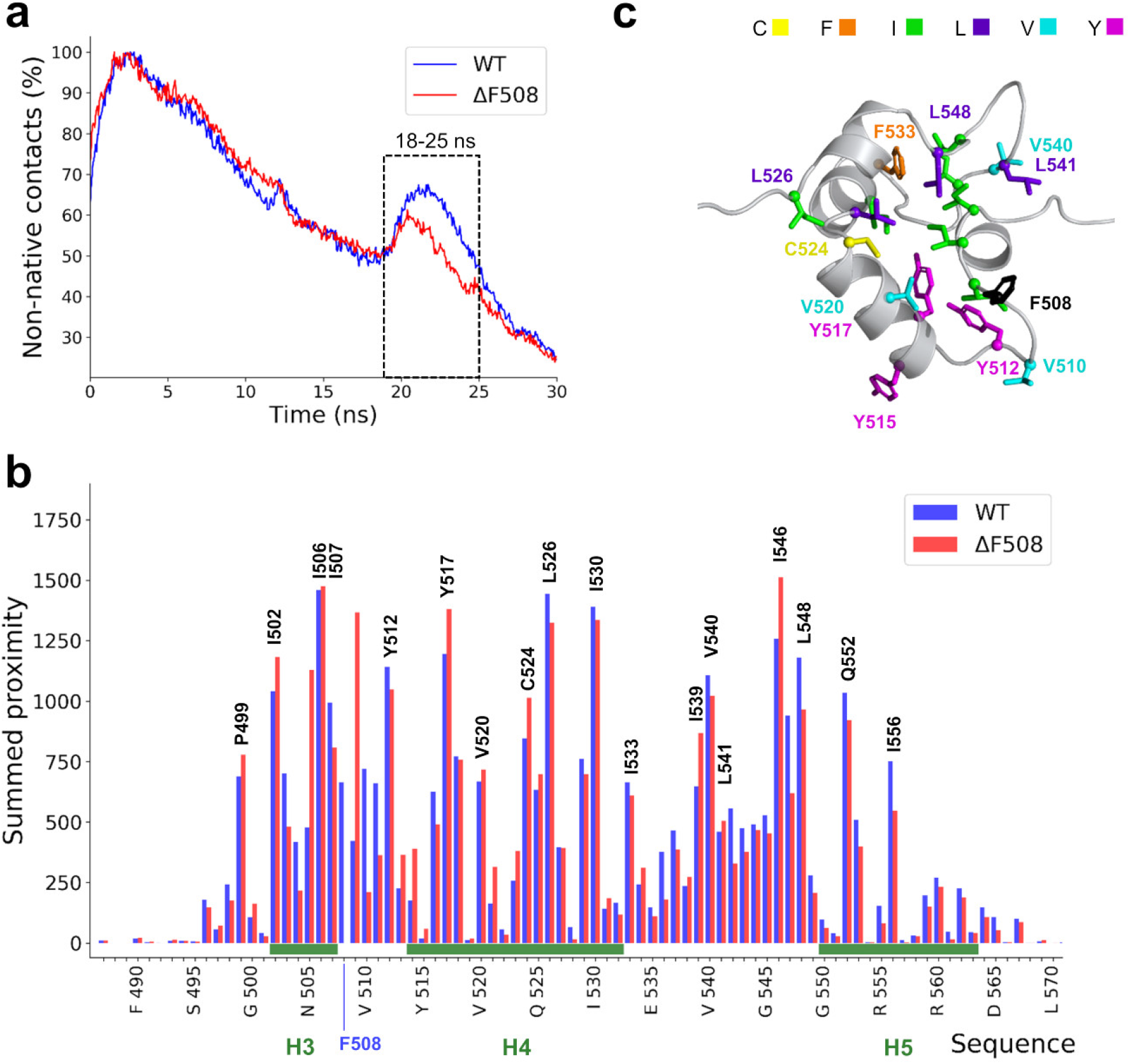
An increased number of non-native contacts in WT may support self-chaperoning. (**a**) The number of non-native contacts normalized to the maximal value during pulling (n=50-50) is plotted. (**b**) Summed proximities of non-native contacts from all trajectories along the sequence of S6-α-S8 core. A higher value indicates that the residue has many close interactions during the investigated time period of unfolding (18-25 ns), which interactions are not present in the native structure. Blue and red columns represent the wild type and ΔF508 mutant, respectively. (**c**) Amino acid positions with the highest proximity values forming a hydrophobic core in the wild type protein are shown on the S6-α-S8 core structure (centroid of cluster #3) by stick representation.

In the next step, we analyzed which residues were involved in the non-native contacts observed in these conformations between 18 and 25 ns (Fig. 4, Supplementary Fig. 9-11). Per residue proximity values were exploited to determine those residues which participated in non-native contact formation (Fig. 4b, and Supplementary Fig. 10-11). Residues exhibiting pronounced differences between the WT and mutant proteins were located in the H3-H4 loop around F508 and in the H4-H5 loop. Most of the residues in non-native contacts were hydrophobic and located in the gap between helices H3, H4 and H5 (Fig. 4c). The above studied non-native intermediate was the last one during unfolding, thus it is likely formed as the first intermediate in the reverse, folding process. This suggests that F508 may play an important role in the interaction network of a non-native intermediate during the early folding that was also observed in our folding simulations (Supplementary Fig. 12). Clustering and contact analysis were also performed on simulations using 0.1 m/s pulling speed (Supplementary Fig. 8, 9 and 10) and indicated some differences from simulations at higher pulling speed. We discussed this in the supplementary materials.

### AFM experiments revealed distinct unfolding steps of NBD1

For the pulling experiments the NBD1 N-terminus was tagged with SUMO (small ubiquitin-like modifier) protein (Fig. 5a). We applied SUMO as a fusion protein, since it has been reported to enhance protein expression and it possesses a well-characterized unfolding pattern in force-extension curves obtained by AFM, aiding selection of successful experiments and their analysis [41]. The C-terminus of NBD1 was cross-linked to the mica-surface and the N-terminal SUMO tag was grabbed and pulled via non-specific binding. A successful pulling event is characterized by typical saw tooth profiles with peaks indicating rupture events, by a total length of ∼110-130 nm (length of the SUMO-NBD1 protein) and a terminal contour length increment of ∼25 nm (the SUMO fingerprint) (Fig. 5b).

**Fig. 5:**
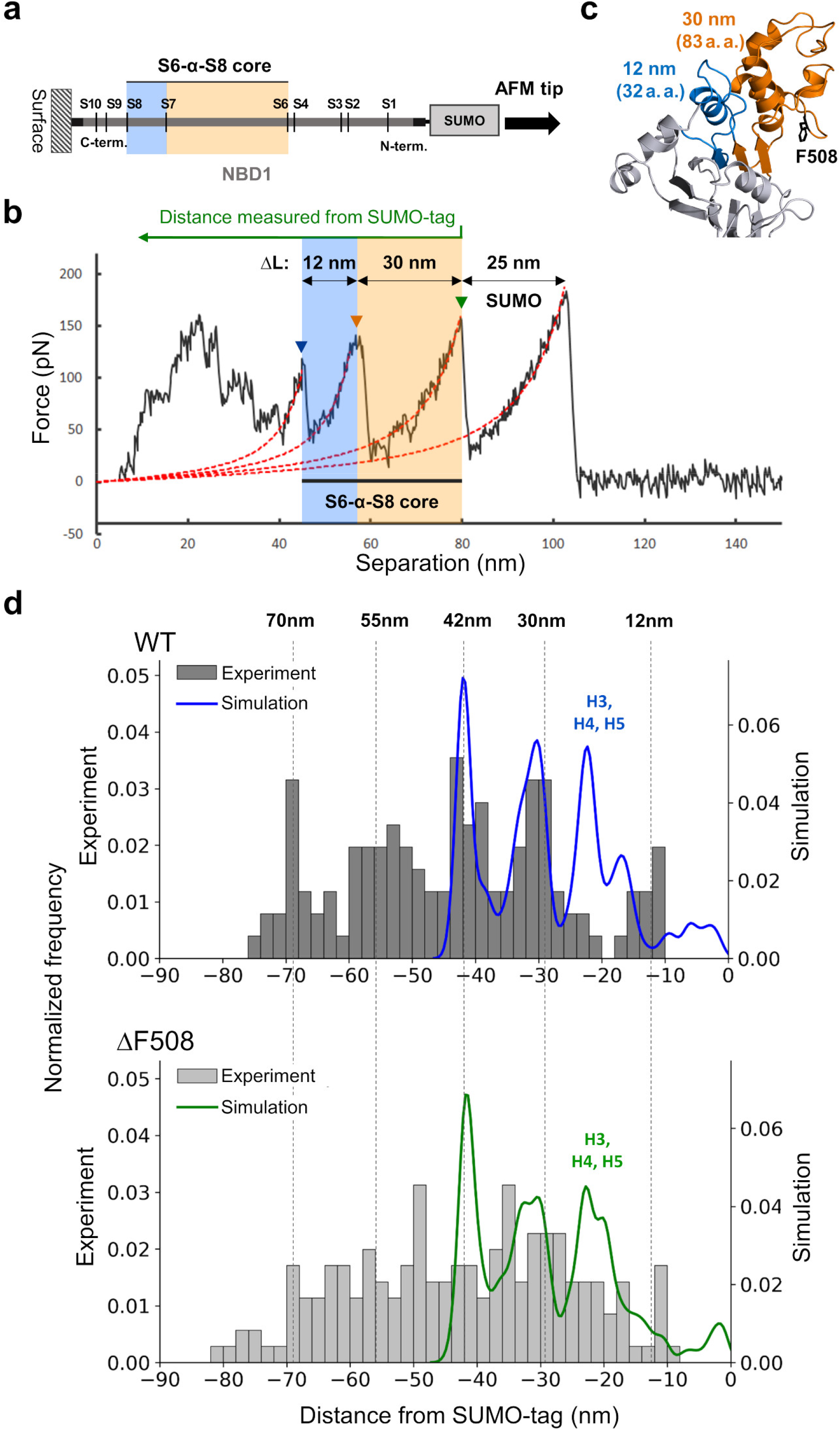
ΔF508 mutant exhibits a dispersed contour length distribution, acquired by AFM, when compared to wild type. (**a**) Schematic illustration of the cysteine-less ΔRI construct. SUMO-tagged NBD1 is immobilized on mica surface via a disulfide bond. The approximate position of β-strands is marked with S letters. (**b**) A force-extension curve demonstrating a characteristic sawtooth profile. Unfolding events are marked by triangles. Peaks are fitted with the worm-like chain model (red dashed lines), providing contour length values (ΔL) from the unfolding of intermediates. (**c**) The structural regions corresponding to the unfolding regions with the given unfolding lengths are colored blue and orange on the NBD1 structure. (**d**) Histograms show the experimental contour length increments measured from the SUMO-tag (WT, n = 126; ΔF508, n = 178). Increments corresponding to the expected length increases based on structure and simulations are indicated with dashed lines. Kernel density estimates (kde) show the distribution of contour length increases from S6-α-S8 core pulling MD simulations (blue -WT, green - ΔF508).

In order to quantitatively compare the pulling events of WT and ΔF508-NBD1 constructs, contour length increments (ΔL) derived from the NBD1 unfolding, detected before the unfolding of SUMO, were collected from force-extension curves. For WT, the ΔL histogram exhibited five peaks (Fig. 5d). In order to correlate these peaks to molecular events, we used the unfolding pathway set of natively folded NBD1 obtained by Gō simulations and NBD1 structural information. In simulations, the last event was the breaking of the S6-α-S8 core. This core consists of two SSE groups, gS8 and gS6. The SSEs were grouped when they unfolded together in Gō simulations (Supplementary Fig. 1). For example, the β-strand S8 and the α-helix H6 together form the group gS8. Since groups gS8 and gS6 are regions of 32 and 83 amino acids in length, respectively, their detachments result in contour length increments of ∼ 12 nm (32 × 0.35 nm) and ∼ 30 nm (83 × 0.35 nm). Depending on the order in which the gS8 and gS6 regions of a given NBD1 unfold, and whether they unfold separately or together, contour length increments of 12, 30 and 42 (12+30) nm resulted during the unfolding. These data corresponded to well-defined peaks observed in the last part of the force-extension curve (Fig. 5). Breaking of group gS3 resulted in the unfolding of a ∼13 nm (36 a.a.) long segment. If this unfolding happened together with the breaking of gS8 and gS6, then the produced contour length increment was 55 (42 + 13) nm. By similar reasoning, breaking of gS10 and gS9 at the start of unfolding produced a 74 or 72 nm peaks in the force-extension curves, respectively. The first two peaks in the histogram of the contour length increments (Fig. 5d, upper histogram) were around 55 and 70 nm. Albeit these data were noisy, since they included tip positions close to the mica surface with adverse electrostatic interactions, the changes in contour length corresponded quite well to these specific breaking events of WT NBD1.

### ΔF508 mutation decreased the proportion of natively folded S6-α-S8 core of NBD1

ΔF508-NBD1 pulling experiments were analyzed as that of WT. The histogram built from ΔF508-NBD1 data did not contain well-defined peaks when pulling WT NBD1 in AFM experiments, but the distribution of contour length changes was homogeneous. Additionally, in a large number of pulling experiments with the ΔF508 mutant, unexpected contour length increments were observed as peaks at approximately 20 and 35 nm compared to WT (Fig. 5d). These peaks suggested that a significant number of ΔF508-NBD1 exhibited modified mechanical resistance or incorrect folding.

In order to resolve these two mechanisms, our MD simulations and experiments were correlated. We calculated contour length increments from force-extension curves of our fully solvated atomistic simulations with the S6-α-S8 core using the WLC model as in experiments. We found that the histogram peaks at 42 nm and 30 nm from MD simulation with WT NBD1 match the experimental data of WT NBD1 suggesting that we were able to detect the gS8 and gS6 unfolding by AFM (Fig. 5d). Since we pulled a natively folded S6-α-S8 core region in our simulations, the similarity of the ΔL *in silico* and *in vitro* peaks confirmed that these were the peaks characterizing the mechanical resistance of the correctly folded structure. Unfolding of α-helices H3, H4 and H5 as force peaks can be observed in the simulations, albeit they did not emerge in experiments likely because of the buffered unfolding of α-helices in *in vitro* experiments [42] (see below, Supplementary Text, and Fig. 5).

The same analysis performed for the ΔF508 simulations resulted in a histogram, which was highly similar to the one from the simulations with the WT S6-α-S8 core (Fig. 5d). This was not unexpected, since only the natively folded and not misfolded NBD1 structures were known and applied in simulations. Importantly, the WT-like peaks were observed in spite of decreased forces (Supplementary Fig. 5) in simulations, suggesting that the difference between WT and ΔF508 experimental pulling curves was mainly not caused by a decreased mechanical resistance of the mutant, but by its decreased folding yield. Counting the number of AFM unfolding events corresponding to the breaking of the natively folded S6-α-S8 core (any combination of groups gS8 with 12 nm and gS6 with 30 nm ΔL) showed that the unfolding signature of the S6-α-S8 core was native-like only in 28% of the ΔF508 experimental curves that is exactly a twofold decrease when compared to the 56% of WT curves with these peaks (Table S1).

### The corrector molecule BIA acts on the α-helical subdomain

A small compound, BIA (5-bromoindole-3-acetic acid) has been demonstrated to promote ΔF508-CFTR maturation and modestly stabilize the ΔF508-NBD1 against thermal unfolding [43]. Therefore, we investigated the effect of BIA on the unfolding of the mutant NBD1 in AFM experiments. We compared the contour length increments of ΔF508-NBD1 in the presence and absence of BIA and found the appearance of a pronounced peak around 24 nm in the presence of this compound (Fig. 6a and Supplementary Fig. 13). The typical WT peaks of 42 and 30 nm were not observed in the presence of BIA, suggesting that this molecule did not restore the native, WT-like conformation or mechanical properties of NBD1, but it bound to and stabilized an NBD1 region, resulting in a peak around 26 nm. This peak is close to the H5-H4-H3 peak observed in simulations, indicating that BIA binds to this part of the NBD1 α-subdomain. This observation also suggests that BIA binding provides improved stabilization of this region than α-helices would alone in the native structure, since with our AFM setup the helix unfolding was not detected in the case of WT (Fig. 5d).

**Fig. 6:**
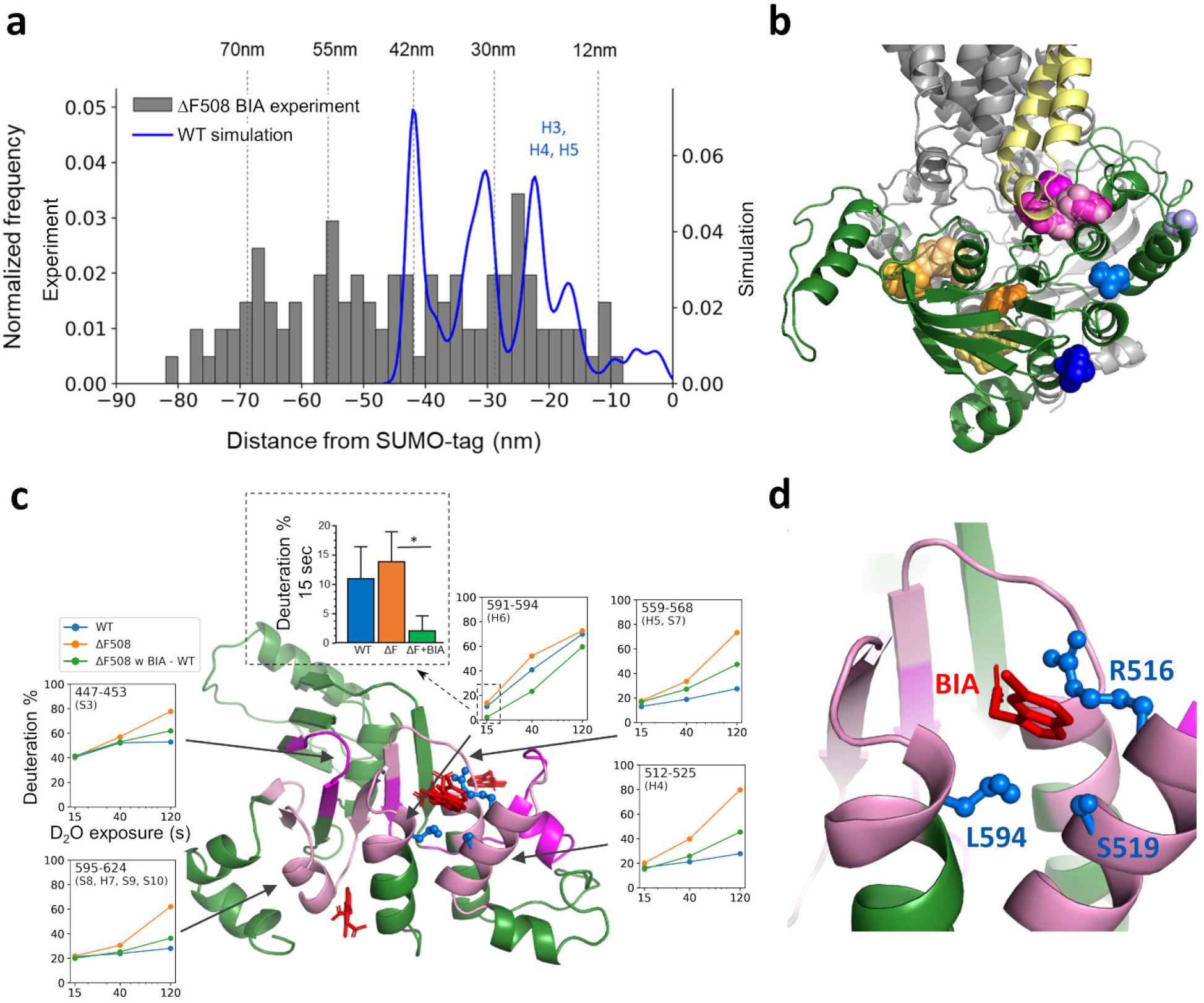
BIA partially restores the WT unfolding pattern and binds to the α-subdomain. (**a**) The histogram shows the experimental contour length increments measured from the SUMO-tag, in the presence of BIA (n = 126). Increments corresponding to the expected length increases based on WT pulling are indicated with dashed lines. Kernel density estimates show the distribution of contour lengths increases from WT S6-α-S8 core pulling MD simulations (blue). (**b**) Potential drug binding pockets in NBD1 were identified by FTMap. Some small molecule fragments bind to CL4 binding pocket (pink spheres) or NBD1/NBD2 interface (orange spheres). Green: NBD1, yellow: CL4, gray: other parts of CFTR. Binding of fragments (blue spheres) to rational pockets does not interfere with CFTR assembly. (**c**) BIA (red sticks) was docked to NBD1 using Autodock Vina. The locations exhibiting higher or lower decrease in HDX are colored pink or magenta, respectively. Corresponding HDX rates for these peptides in WT- (blue) and in ΔF508-NBD1 in the absence (orange) or presence (green) of BIA are shown as insets. The significant BIA inhibition on the ΔF508-NBD1 a.a.591-594 peptide deuteration is depicted by the bar plot. (**d**) The docked pose with the best binding score indicates BIA interaction with L594 beside R516 and S519 of the ΔF508-NBD1.

Two peaks around 70 nm and 55 nm were also detected in the presence of BIA, which likely correspond to the β-subdomain and were observed in WT but not in ΔF508. Thus, we aimed to narrow the potential BIA binding sites using *in silico* methods. FTMap^44^ identified several putative drug binding sites in NBD1 (Fig. 6b), but sites at NBD1/CL4 and NBD1/NBD2 interfaces were excluded, since binding to those locations would interfere with CFTR assembly and maturation. Two out of three sites in the α-subdomain were reinforced by Autodock Vina [44] docking (Fig. 6c and Supplementary Fig. 14). One includes ends of H4, H5, H6 and loops, and another involves H6, the loop between S2 and S3, and H7.

Additional evidence for the BIA ligand-binding site was sought for by determining the backbone amide hydrogen deuteration of the isolated ΔF508-NBD1 in the absence and presence of BIA using the hydrogen deuterium exchange and mass spectrometry (HDX-MS) technique. The time course of the domain deuteration was monitored by continuous labeling for 15, 40 and 120 sec at 37°C. The accelerated deuteration of the ΔF508-NBD1 relative to the WT-NBD1 was consistent with thermal destabilization of domain that was reported previously [13, 16, 45]. The ΔF508-NBD1 HDX was partially suppressed at four α-subdomain peptides, including of H4 (a.a. 512-525), H5 (a.a. 559-568) and H6 (a.a. 591-594), and at a group with several secondary structure elements (end of H6, S8, H7, S9 and S10; a.a. 595-624) (Fig. 6c). Importantly, BIA was able to attenuate the HDX kinetics of the mutant below the WT-NBD1 in a single peptide of the H6 (a.a. 591-594) at all incubation times. According to in *in silico* docking calculations, the last residue of the H6 (a.a. 591-594) peptide, L594 sidechain serves as a docking surface to BIA (Fig. 6d). These observations strongly suggest that BIA bound this region of H6 and its stabilizing effect is communicated allosterically to distant parts of NBD1, also explaining the native-like AFM unfolding signature of ΔF508-NBD1 β-subdomain in the presence of BIA.

## Discussion

In this study, we explored the mechanical unfolding of CFTR NBD1 domain using both atomistic molecular dynamics (MD) simulations and single molecule force spectroscopy (AFM) experiments. We found that deletion of F508 has a significant effect on NBD1 unfolding. Our results suggest that these effects were likely exerted by hindering the formation of crucial non-native intermediate states in the late stages of unfolding, thus potentially in the early stage of folding.

Although the first step of CFTR biogenesis, which is affected by ΔF508, is the folding of NBD1, there are only a handful of studies investigating this process. Qu *et al*. demonstrated that ΔF508 increased the probability of off-pathway intermediates and affected an early folding step before the formation of the ATP binding site [22, 23]. They used isolated NBD1 and measured its folding yield by light scattering and intrinsic Trp fluorescence, showing that F508 affects the rate of maturation and suggested that F508 makes crucial contacts during the folding process. Interestingly, it has also been demonstrated that deletion of the three nucleotides resulting in ΔF508 mutation causes alteration in mRNA structure, leading to a decreased rate of translation [24]. The authors also described that the lower translation speed resulted in altered CFTR conformations in metabolic pulse chase experiments [25]. A set of other experiments targeted NBD1 folding on ribosomes using truncation constructs and FRET [17, 26, 27]. These studies revealed that the folding of the N-terminal part involving β-strands S1-S6 took place while F508 was in the ribosome tunnel and the deletion affected a later stage of domain folding [17]. The authors found that the ribosome delayed the α-subdomain folding which was essential in the correct insertion of S7/S8 strands into the β-sheet core [26]. These results seem to be contradictory to that of Qu *et al*. [22, 23]. However, the N-terminal β-subdomain may fold rapidly and independently according to FRET experiments by Kim *et al*. [26], but the state of this subdomain is likely not sufficiently mature to provide an environment for forming a correct ATP binding site [22, 23]. Most likely the different levels or timescales of experiments provided data on different aspects of the folding process. Although the above studies on NBD1 folding were fundamental and agreed in the involvement of altered intermediate states, their resolution was low.

Due to the success of using simplified models in pulling simulations [46, 47], we first used a native structure based Gō model that overcomes computational limitations associated with pulling simulations with NBD1-sized proteins. This simplified model was essential for the analysis of the force-extension curves. However, because its accuracy is limited by the lack of explicit water molecules and non-native contacts, we also performed regular, fully solvated atomistic force field pulling simulations with a smaller part of NBD1, the S6-α-S8 core. A detailed analysis of the unfolding was performed on the results of these simulations. We detected altered pathway frequencies and faster detachment of certain secondary structure elements in the mutant core (Fig. 2 and Supplementary Fig. 3), suggesting differences in the interaction network around the β-strand S6 and in the final unfolding intermediate unit containing α-helices H4, H5 and H3. Importantly, our results showed that F508 remained a component of the folded part of the wild type NBD1 almost until the end of unfolding (Fig. 3 and 4). The WT core exhibited a higher number of non-native interactions at a late stage of unfolding compared to ΔF508, suggesting that non-native interactions contribute to the stability of the late unfolding intermediate detected in the wild type core. These interactions included positions with known CF-causing mutations (ΔI507, V520F, L558S and A559T) that have been shown to affect the α-core compaction of the nascent NBD1 during a critical window of folding (Shishido *et al*. [27]).

Taken together, F508 supports the development and persistence of non-native interactions that may be an important factor for off-pathway avoidance and self-chaperoning. The non-native contacts, which have been described to influence the folding free-energy barrier [48] and can become the rate-limiting step of protein folding [49], likely serve as a deceleration mechanism to provide time for the NBD1 polypeptide to acquire the right intermediate state before engaging the next step of folding. This was also supported by the *in vitro* translation experiments of Kim *et al*., showing that faster codons inhibited folding [26]. The same residues, which were involved in the formation of non-native contacts during unfolding (Fig. 4 and Supplementary Fig. 10), were also in contact in our folding simulations (Supplementary Fig. 12), confirming that the residue F508 and its surroundings may serve as a folding nucleus. Earlier, a decreased folding time was observed for the ΔF508-NBD1 in Gō folding simulations, suggesting that the self-chaperoning of NBD1 was diminished [50].

Importantly, by combining experiments and computer simulations, we identified drug binding sites that are located on the surface of NBD1 away from the protein axis and exposed to the solvent. Therefore, drug binding to these regions is unlikely to interfere with CFTR domain-domain assembly and maturation (Fig. 6b, c). In addition, drugs targeting these regions may not only rescue the volatile folding and stability of the α-subdomain, but potentially allosterically stabilize the β-subdomain, as confirmed experimentally by BIA binding (Fig. 6a, d). We also demonstrated by computational methods that secondary site mutations either in the β-subdomain or in the α-subdomain restored the WT-like allosteric network in the absence of F508 (Supplementary Fig. 16). Because of this allosteric subdomain coupling, we propose that a drug rationally designed to bind the α-subdomain, not only corrects ΔF508 and other mutations in the α-subdomain, but also has the potential to rescue CF mutations localized in the β-subdomain.

In summary, we found that the deletion of F508 has a significant effect on the unfolding pathways of NBD1 and accelerated the detachments of certain secondary structure elements compared to the wild type. Our results suggest that these effects were likely exerted by hindering the formation of crucial non-native intermediate states in the late stages of unfolding, thus potentially in the early stage of folding (Fig. 3-4 and Supplementary Fig. 8-11). The experimental results suggest that the S6-α-S8 core is folded incorrectly in a significant portion of the wild type NBD1, and that misfolding is greatly enhanced by the F508 deletion. We conclude that the α-subdomain has an inherited property for folding instability. We propose that the instability-enhancing effect of NBD1 mutations may be corrected by small molecules binding to the α-subdomain, allosterically stabilizing the full domain. Furthermore, our results confirm that F508 maintains a network of non-native contacts and suggest a role in slowing down the translation, thereby aiding self-chaperoning.

## Methods

### Structural models

Wild type NBD1 structure based on an X-ray structure (PDBID: 2BBO) from an earlier study [51] was used as the starting point. In order to match the construct used in our experiments, regulatory insertion (a.a. 405-435) was removed and the gap was sealed by loop modeling of Modeller [52], setting residues 403, 404, 433, and 434 as a loop region. ΔF508 mutation was modeled similarly. The missense mutations were generated using the mutagenesis tool of VMD [53].

### Molecular dynamics simulations

Conventional all-atom MD simulations were performed with the WT and ΔF508 S6-α-S8 core region of the NBD1 structure. The S6-α-S8 core (a.a. 487-604) consists of three β-sheets (S8, S7, S6) and five α-helices (H3, H4, H4b, H5, H6), including F508. MD simulations were run using GROMACS 2019 [54] with the CHARMM36m [55] force field and the TIP3P water model. A 150 mM KCl concentration was used. Hydrogen atoms were replaced with virtual interaction sites to speed up the calculations, such that a 4 fs time step could be used [56]. Electrostatic interactions were calculated using the fast smooth PME algorithm [57]; the LINCS algorithm [58] was used to constrain bond lengths.

All structures were energy minimized using the steepest descent integrator in the first step, then equilibration (NVT, NPT) procedure was performed prior to each pulling simulation to generate inputs for independent simulations with different starting velocities (T=310 K, p=1 bar). The Nose– Hoover thermostat and the Parrinello-Rahman barostat with isotropic coupling were employed for the production runs. Time constants for the thermostat and the barostat were set to 2 picoseconds and 5 picoseconds, respectively. The C-terminus of the S6-α-S8 core structure was restrained, and the N-terminus was subjected to constant velocity pulling. Two pulling velocities, 1 m/s and 0.1 m/s were used, requiring 40 ns and 400 ns long trajectories, respectively. 50 simulations were performed with both constructs and for each of the two pulling speeds. MD parameter files can be downloaded from http://resources.hegelab.org.

### Protein expression and purification

We used a cysteine-less [59], His6-tagged SUMO-fusion NBD1 carrying a deletion of regulatory insertion (ΔRI), which improves the protein stability and solubility [15]. It was especially important for the ΔF508 mutant [15, 16, 41]. Cysteine-less construct was used to avoid interfering with immobilization via terminally introduced cysteines. For simplicity, we referred to Cys-less NBD1 ΔRI construct as NBD1. SUMO-NBD1 constructs were purified from *E. coli* Rosetta 2 (DE3) pLysS strain. His6-tagged proteins were purified using an Ni-NTA affinity column (Profinity IMAC Ni-Charged, BioRad).

### Single-molecule force spectroscopy experiments and analysis

Freshly cleaved mica surface was functionalized with APTES. The terminal cysteine of NBD1 was cross-linked to the APTES-coated mica using Sulfo-SMCC. Force spectroscopy was carried out on a Cypher atomic force microscope (AFM) instrument (Asylum Research) using PNP-TR cantilevers (spring constant: 100-200 pN/nm, NanoWorld). Experiments were performed at 25 °C in PBS buffer (pH 7.2) [60]. Unfolding of NBD1-SUMO was carried out by first attaching the protein to the tip non-specifically by applying a constant force of 1 nN for 1 s to the tip on the mica surface, then followed by constant speed retraction with a pulling velocity of (1 μm/s).

Data was fitted using the worm-like chain (WLC) model of polymer elasticity using Igor Pro (Wavemetrics) extended with the Asylum Research AFM driving software. Since AFM tip and protein adhesion occurred at random locations, most of the retraction curves did not show the unfolding of the full protein. Force-extension curves exhibiting an overall length compatible with a completely unfolded NBD1-SUMO protein (total length of ∼110-120 nm) and including the SUMO unfolding fingerprint as a terminal contour length (Lc) increase of ∼25 nm were selected. All calculated Lc values were collected and summarized in histograms.

### Analysis of unfolding pathways

The analysis was completed using GROMACS tools [54], the MDAnalysis package [61] and in-house Python scripts. For identification of secondary structural units’ detachment over the pulling trajectories, the native contacts (contacts in the initial folded structure) of every secondary structural unit were determined and the fraction of native contacts (Q) was calculated as a function of time over each pulling trajectory using the *soft_cut* metric from the MDAnalysis package [61, 62]. We defined a detachment event if Q decreased below a threshold value, which was set 0.2 in the case of Gō simulations with NBD1 and 0.1 in the case of conventional MD simulations with the S6-α-S8 core region. The structures of the unfolding simulations were clustered to identify intermediate states during unfolding. We used a pairwise contact-based RMSD with a cutoff value of 0.8 nm as a distance metric for clustering as described by Mercadante *et al*. [40, 63]. The pairwise residue distance matrix of all unfolding conformations along every pulling trajectory was calculated and all values above the 0.8 nm cutoff value were set to 0.8 nm. The pairwise contact-based root-mean-square deviation (RMSD) values were calculated from these distance matrices, as

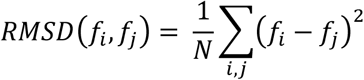

where N is the number of residues and f_i_ is the pairwise a.a. distance matrix of a given frame with position index i. Using this type of RMSD values for clustering led to ignoring the changes between distant residues that would have masked changes in important contacts within the remaining folded part. We applied a density-based clustering algorithm, DBSCAN for clustering [64]. The structures of each pulling simulation were clustered separately using the contact-based RMSD as a distance measure and the centroids of these clusters were pooled. The centroid structures from all simulations were clustered again to yield all observable intermediates from every simulation. Clusters with fewer members than 5% of the clustered structures were omitted from the re-clustering step. The DBSCAN clustering parameter, Eps, was set based on the Elbow method [64] and considering the Silhouette Score [65]. The cluster centroids were selected based on a calculated similarity score [66] as

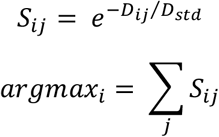

where D: pairwise distances of the structures of the cluster (contact based RMSD values), S_ij_: pairwise similarity of two structures, D_ij_ (RMSD_ij_): pairwise distance of structures i and j from the conformational ensemble (cluster), D_std_: standard deviation of D, argmax: the structure most similar to all other structures within the cluster (centroid).

### Proximity calculations

The number of native and non-native contacts during each unfolding trajectory were calculated and normalized. Amino acid residues were in contact if the distance between any atom of the two residues is smaller than the 0.45 nm cutoff value. Contacts between residues that were present in 75% of the initial structure of the 50 simulations were labeled as native. Residues that were not native contacts but got closer than the cutoff in the course of the unfolding simulations were assigned as non-native contacts. Proximity values [67] were used to identify important non-native contacts where their cumulative number was increased during the simulations (18-25 ns and 180-250 ns range at pulling velocities of 1 m/s and 0.1 m/s, respectively). Proximity is minimal (zero) for the cutoff value or larger distances and maximal (one) if the distance between two residues is zero:

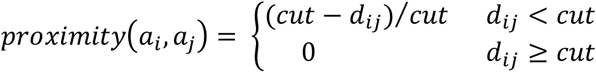

where cut: distance cutoff between amino acids (0.45 nm), a_i_: amino acid residue in position i, d_ij_: distance between amino acids in position i and j. Proximity values for amino acid pairs in each structure were summed for all the WT and the ΔF508 simulations (separately for both pulling velocities) in the time interval of interest (Supplementary Fig. 9). The summed 2D proximity values for each amino acid residue were also calculated by summing the previously calculated proximity values of the contacts of the given residue (Fig. 4 and Supplementary Fig. 10).

### Calculation of rupture forces and contour length increments in simulations

Rupture forces were derived from the pulling trajectories. Peaks of the force curves were collected for each SSE around the detachment time point of the given SSE. Normalized frequency of rupture forces from all and from individual SSE detachment events and their Gaussian density estimates were calculated and visualized. Contour lengths and their increments (ΔL) were calculated using the simple polynomial worm-like chain (WLC) interpolation formula [68].

### Identification of drug binding sites in NBD1

FTMap webserver[69] (http://ftmap.bu.edu/) was used with default options to identify potential binding pockets on the NBD1 surface. Docking of BIA was performed with Autodock Vina [44]. Default options were applied except exhaustiveness, which was increased to 128. The search space was defined as described and shown in Supplementary Fig. 14.

### Visualization

Structures are visualized using PyMOL (The PyMOL Molecular Graphics System, Version 1.8.4 Schrödinger, LLC). Figures were generated by Matplotlib [70].

### Hydrogen-deuterium exchange (HDX) experiments

The ΔF508-NBD1 of human CFTR was purified as described [13]. Deuteriation time course of the ΔF508-NBD1 was measured by HDX coupled with mass spectrometry (HDX-MS) technique [71]. The sample concentration was 5 µM in buffer containing: 10 mM HEPES, 150 mM NaCl, 1 mM ATP, 2 mM MgCl_2_ and 1 mM TCEP at pH 7.5. The deuterium uptake was performed in D_2_O-based buffer in the presence and absence of 2 mM BIA. For each deuteration time, NBD1 was mixed with 1:14 dilution ratio into D_2_O-based buffer, resulting more than 90% D_2_O contents, and incubated for 10 s, 40 s, and 120 s. HDX reaction was quenched by adding chilled quenching buffer (300 mM glycine and 8 M urea at pH 2.4) with 1:2 ratio. Quenched solution was flash frozen in MeOH containing dry ice and stored at -80 °C until use. 10 μL of quenched sample was injected into the sample loop, followed by in an on-line immobilized pepsin column prepared in house. On-line pepsin digestion was carried out at a flow rate of 50 µL/min for 1.5 min, and resulting peptides were trapped on a C18 trapping column (Optimized technologies, Oregon City, OR). Following desalting for 1.5 min at a flow rate of 180 µL/min, the peptides were loaded onto a C8 analytical column (1 mm i.d. × 50 mm length, Thermo Fisher Scientific) and separated with Agilent 1290 Infinity II UHPLC system. Separated peptides were detected by LTQ Orbitrap XL (Thermo Fisher Scientific) in positive-ion mode for m/z 200 – 2000 using electrospray ionization. For peptide identification, tandem MS (MS/MS) analysis in data-dependent acquisition mode with collision-induced dissociation was performed in separate measurements. All MS/MS data were analyzed in Proteome Discoverer 1.4 (Thermo Fisher Scientific). The deuteration were determined from triplicate measurements and the collected data were analyzed using HDExaminer 2.3 (Sierra Analytics). The relative deuterium uptake (%D) for each peptide was calculated by comparing the centroids of the isotope envelopes of the deuterated samples against the undeuterated controls. Deuterium uptake plots were generated using Prism 6 (Graphpad).

## Acknowledgements

We are grateful to K. Lór for excellent technical assistance. We thank M. Habibi and S. Plotkin (University of British Columbia, Canada) for their help in setting up simplified pulling simulations. We acknowledge the computational resources made available on the GPU cluster of the Governmental Information-Technology Development Agency (https://kifu.gov.hu), the Grubmüller laboratory at Max Planck Institute (https://www.mpibpc.mpg.de/grubmueller), and Wigner GPU Laboratory (http://gpu.wigner.mta.hu).

## Data Availability

All input data are available from the authors upon request. **Code availability**. Scripts are available from the authors upon request.

## Author contributions

TH, HG, MK, and GLL conceived ideas; RP, BF, TH, HG, LG, and MK wrote the manuscript, RP, BK, and HT performed the pulling experiments; NS performed HDX experiments; BF and TH performed simulations and their analysis.

