## Supplementary Material for "Nanomechanics combined with HDX reveal allosteric drug binding sites of CFTR NBD1"

### Supplementary Methods

MD simulations with SMOG2. To perform the mechanical unfolding of NBD1, steered pulling molecular dynamics simulations were performed using the all-atom Gō model [1] of the SMOG2 tool [2]. Input files were generated for GROMACS 4.6.7 using SMOG2 tools (e.g. *smog_adjustPDB* with default parameters and *smog2*). Options for neighbour searching, non-bonded interactions, generalized Born electrostatics, pressure and temperature coupling were set according to SMOG defaults and parameters used by Habibi *et al.* [3]. Temperature was set to 110K, which is around the melting temperature of an AA-SMOG system of NBD1 size. 100 simulations were initiated with different initial velocities for WT NBD1 construct. In the 1 ns long equilibration step back-bone and side-chain atoms were restrained using 400 and 40 kJ/mol/nm$^2$ harmonic force constants, respectively. The C-terminal NBD1 end was restrained, and the N-terminus was exposed to constant velocity pulling (5 m/s) for 16 ns using GROMACS pull code to achieve a fully stretched polypeptide at the end of the simulations.

Folding simulations. In order to test the initial steps of the NBD1 α-subdomain folding, replica exchange (REX) discrete molecular dynamics (DMD) simulations were performed [4]. The involved all-atom model of the polypeptide with the CHARMM-based Medusa force field and implicit water model made the folding simulation of a.a. 491-567 feasible. Elongated peptide structure was generated by a CHARMM script and used as input. Eight replicas at temperatures 0.5246, 0.5451, 0.5665, 0.5886, 0.6116, 0.6355, 0.6604, and 0.6862 temperature unit were run for 1,000,000 time unit. Conditions for replica exchange were tested every 1,000 time units and these temperatures provided 23-51% exchange probability between neighboring replicas. Frames were saved every 200 time units, thus 5,000 conformations were generated for each temperature. Anderson's thermostat was used, and the heat exchange factor was set to 0.01. Frames from every trajectory were grouped by temperature and the conformations at the highest temperature were used for analysis.

Equilibrium simulations and their network analysis. NBD1 structure generated based on an X-ray structure (PDBID: 2BBO) was used as the starting point [5]. ΔF508 mutation was modeled using the loop modeling algorithm of Modeller [6]. The I539T rescue and other second site suppressor (1S [F494N] and 3S [F494N, F429S, and Q637R]) constructs were generated using VMD [7]. Simulations with these constructs were run to match our early HDX and AFM experiments with NBD1 in the 1S background. However, finally we were able to purify NBD1 for these experiments in the absence of suppressor mutations. Although suppressor mutations tend to reduce the difference in the WT- and ΔF508-NBD1 conformational stability, we obtained meaningful differences in MD simulations in the presence of 1S suppressor background (which account for an ~1.5°C increase in the ΔF508-NBD1 meting temperature) [8], thus our findings can be considered valid. Equilibrium simulations were prepared by CHARMM-GUI [9] as follows. Terminal residues were patched by ACE (acetylated N-terminus) and CT3 (N-methylamide C-terminus), 150 mM KCl in TIP3 water were used, grid information for PME (Particle-Mesh Ewald) electrostatics was generated automatically, and a temperature of 310 K was set. The structures were energy minimized using the steepest descent integrator (maximum number to integrate: 50,000 or converged when force is <1,000 kJ/mol/nm). From the energy minimized structures parallel NVT equilibrium simulations were forked, followed by production runs for 0.5 μs. Nose-Hoover thermostat and Parrinello-Rahman barostat were applied. Electrostatic interactions were calculated using the fast smooth PME [10] algorithm, and LINCS [11] algorithm was used to constrain bonds. Constant particle number, pressure, and temperature ensembles with a time step of 2 fs were used. Simulations were performed using GROMACS 2018 [12] and CHARMM36 [13] force field. Frames between 450 and 500 ns were selected from every trajectory for network analysis [14], merged, and aligned. Correlated motions in each merged trajectory were calculated based on mutual information [15], using the Wordom package [16]. A network was created by setting residues as nodes and generalized pairwise correlation ($C_{ij}$) was used to set edge weights. Since $0 <= C_{ij} <=1$, the weight was defined as -log(Cij) to have a small distance between highly coupled residues. In order to detect network communities, we used the Girvan-Newman algorithm implemented in the networkx Python package.

### Supplementary Results

*Unfolding pathways of WT NBD1 determined by all-atom Gō simulations.* Gō simulations were performed to determine the NBD1 unfolding pathways and fractions of native contacts between secondary structural elements were calculated. When a structural unit unfolded from the domain, its fraction of native contacts decreased from 1 and approached zero at the time point when its intra-domain contacts were lost due to the applied pulling force (Supplementary Fig. 1). The contacts of usually two or more secondary structure elements diminished together, thus most structural units formed unfolding groups. We marked each unfolding group with one of its structural elements (e.g. the group of S1, S2 and S4 was marked with gS1). The only exception is the β-strand 10 (S10), which segment started unfolding individually.

We determined the sequence of detachments of the groups in each simulation and calculated the frequency of each pathway (Supplementary Fig. 2). Although different pathways could be observed, there were general features. The unfolding of NBD1 always began with the unfolding of the β-sheet subdomain followed by the α-helical subdomain. The detachment of β-strand S10 was followed by the unfolding groups gS9, gS1, and gS3 (Supplementary Fig. 2). The remaining intermediate structure was composed of three β-strands (S6, S7 and S8) and the α-subdomain, which region is referred to as S6-α-S8 core and exhibited two main unfolding pathways. This region was disrupted either by breaking up the S8 element first then the remaining part of NBD1 or by unfolding the whole S6-α-S8 core region at once. Alternative pathways were also observed and usually caused by unfolding groups detaching together at the same time (e.g. groups gS1 and gS9). Two main unfolding pathways were observed for WT NBD1 and several alternative pathways with low probability were also detected (Supplementary Fig. 2).

*Residue contacts in early steps of folding.* The non-native interactions observed in the last WT S6-α-S8 core unfolding intermediate may play a role during the early folding events of this region. In order to test this hypothesis, we performed discrete molecular dynamics folding simulations using replica exchange initiated from a fully stretched peptide corresponding to a.a. 491-567. Since the structures belonging to the highest temperature exhibited the least contacts and probably represented conformations of early folding steps, we used the conformations visiting the highest temperature for analysis.

The region with the highest number of contacts in this set of structures was the neighbourhood of F508 (Supplementary Fig. 12). Four amino acids exhibited a high contact number (F508, Y512, Y515, Y517). Three out of these residues (F508, Y512, Y517) were identified in our unfolding simulations to form non-native interactions. Numerous other amino acids were also found to be relevant as both non-native contacts in the unfolding simulations (Fig. 4b, c and Supplementary Fig. 10) and interaction sites in the folding simulations (Supplementary Fig. 12). In the case of the ΔF508 mutant, the interaction frequencies were slightly increased around a.a. 540-550, which forms the loop (S6c, L2) between helices H4b and H5 in the native structure.

In the neighborhood of the four main interacting residues, the bending of the peptide chain and the formation of an α-helical structure corresponding to the helix H4 was detected during the folding simulation (Supplementary Fig. 12). This region was structurally similar to the last intermediate conformation in unfolding simulations with a major difference, which was the orientation of Y515 side chain. Y515 formed interactions in the folding intermediate but did not take part in the non-native interactions during unfolding. The most frequent contact in the folding simulation was of Y515-Y512, followed by the Y512-F508 contact. We suspect that the lack of F508 side chain disables the formation of an important intermediate structure with hydrophobic contacts of F508, Y512, and Y515.

*Comparison of the WT force-extension curves obtained in AFM experiments and MD simulations.* To further compare the MD simulation and experimental results we evaluated the force-extension curves of S6-α-S8 core fully solvated atomistic MD simulations in the same way as in the AFM experiments using the WLC model. The distribution of contour lengths increases (Kernel density estimates (kde)) from S6-α-S8 core pulling simulations was plotted onto the histograms of contour length increases measured in AFM (Fig. 5). In the case of wild type data, the 42 nm and 30 nm peaks observed in AFM agree well with the peaks of the simulation data. From simulations we know that these events marked the breaking of the interactions between the strands S7-S8 and S6-S7 verifying our presumption based on NBD1 topology. The difference between the experimental and simulation distributions is the location of the last peak. In AFM experiments, the last shorter section, which unfolded, is ~12 nm long, presumably corresponding to the S7-S8 section. Contrarily, in simulations the last event is the breaking of the interactions around the helices H5, H4 and H3 resulting in a strong peak around 22 nm. The lack of this peak in AFM could be caused by the different pulling speed used in experiments and simulations. In AFM we pulled the molecule six orders of magnitude slower resulting in lower loading rates. It is known that the behavior of helices during pulling is complex, and they can act as force buffers under mechanical stress [17]. We presumed that at lower pulling velocity the mechanical resistance of helices is negligible not resulting in considerable peaks on force-extension curves.

*Differences between the formation of non-native contacts in the 1 m/s and 0.1 m/s pulling simulations.* We observed interesting differences between the non-native contact formation of fast (1m/s) and slow (0.1 m/s) pulling simulations. Analyzing the sum of the non-native contacts during the pulling shows that the wild type NBD1 and ΔF508 mutant differ only at fast pulling velocities (Supplementary Fig. 8e, f). To find the reason for this, we analyzed the non-native interactions between the individual amino acids using proximity matrices (Supplementary Fig. 9). At slow pulling velocity the proximity matrices show several, rather compact patches and between them poorly populated areas, which means that the amino acids forming non-native contacts tend to localize in close vicinity to each other (Supplementary Fig. 9 white circled areas). In contrast, at fast pulling velocity the non-native contacts are more scattered. E.g. the surroundings of F508 interact with numerous amino acids along the region of amino acids 508 to 560 (Supplementary Fig. 9 green circled areas). At slow pulling velocity, between the folding steps, the system has more time to equilibrate and reach more localized interacting areas. However, at fast pulling velocity there is less possibility for that. It seems that the differences in the non-native contact formation of wild-type and ΔF508 mutant could be detected only under such conditions. On the other hand, it shows that F508 is able to form interactions with other amino acids under such a dynamic condition.

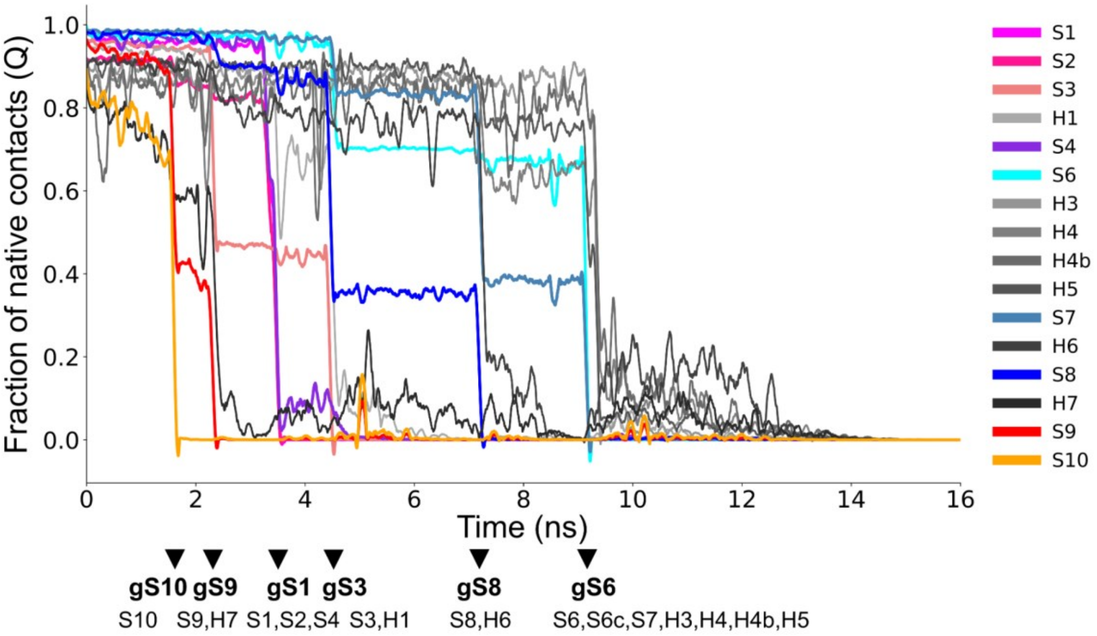


**Supplementary Fig. 1: Unfolding of individual secondary structure units.** Detection of unfolding of secondary structure units in an all-atom Gō simulation as an example. The fraction of native contacts (Q) was calculated for each structural element and plotted. Colored lines: Q of β-strands; gray lines: Q of α-helices; arrows: unfolding events, which are marked with the name of a β-strand in the unfolding group. Unfolding group names and their members are indicated with bold and regular fonts, respectively.

| | | | | | | WT NBD1 |
|---|---|---|---|---|---|---|
| gS10 | gS9 | gS1 | gS3 | gS8 | gS6 | 47% |
| gS10 | gS9 | gS1 | gS3 | gS8-gS6 | | 17% |
| gS10 | gS9 | gS1 | gS3 | gS6 | gS8 | 1% |
| gS10-gS9 | | gS1 | gS3 | gS8 | gS6 | 1% |
| gS10 | gS1 | gS9 | gS3 | gS8 | gS6 | 2% |
| gS10 | gS1 | gS9 | gS3 | gS8-gS6 | | 4% |
| gS10 | gS1-gS9 | | gS3 | gS8 | gS6 | 1% |
| gS10 | gS1-gS9 | | gS3 | gS8-gS6 | | 2% |
| gS10 | gS1 | gS9-gS3 | | gS8 | gS6 | 4% |
| gS1 | gS10 | gS9 | gS3 | gS8 | gS6 | 5% |
| gS1 | gS10 | gS9 | gS3 | gS8-gS6 | | 2% |
| gS1 | gS10 | gS9 | gS3 | gS6 | gS8 | 3% |
| gS1 | gS10 | gS9-gS3 | | gS8 | gS6 | 8% |
| gS1 | gS10 | gS9-gS3 | | gS8-gS6 | | 3% |

**Supplementary Fig. 2: The frequency of alternative unfolding pathways in all-atom Gō simulations using SMOG.**

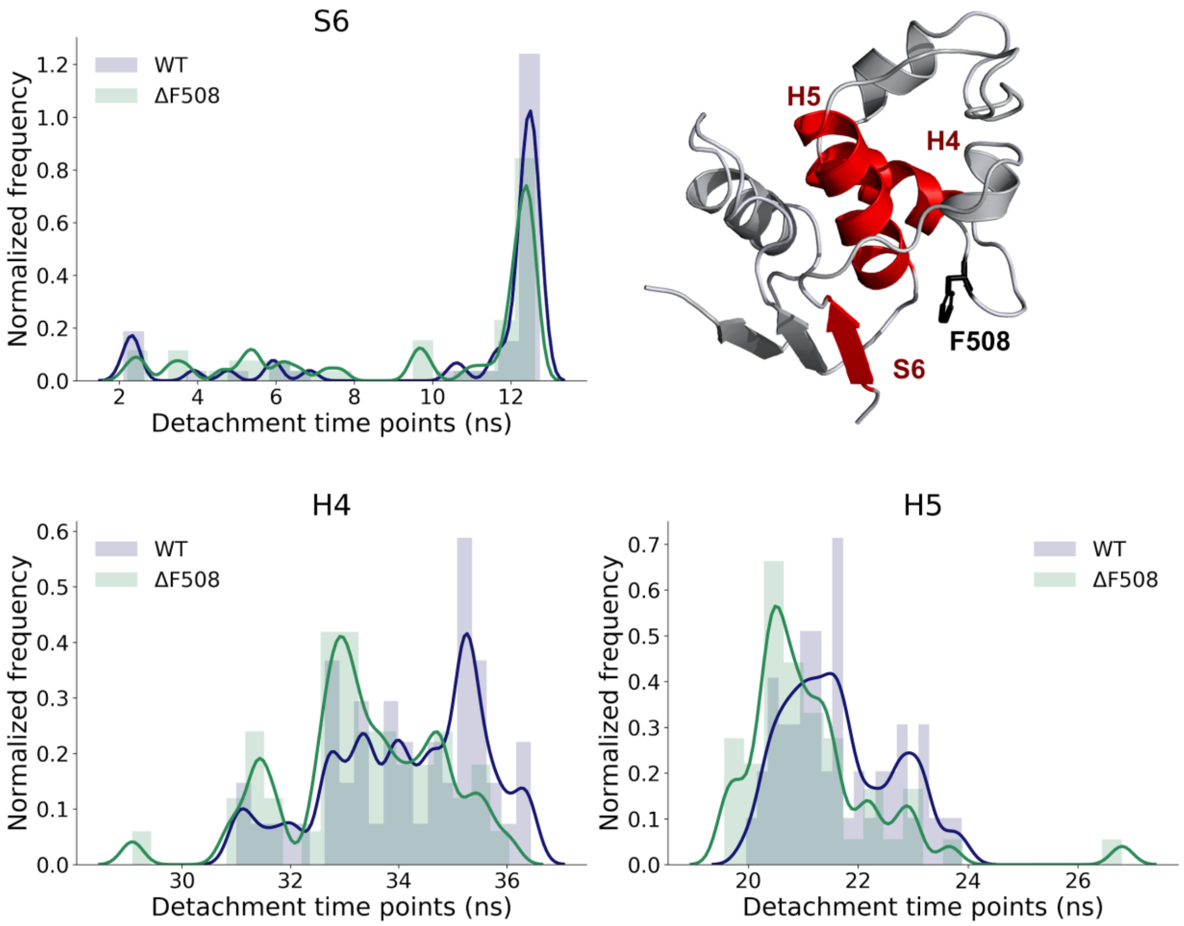


**Supplementary Fig. 3: ΔF508 accelerates the decoupling of specific secondary structure elements in the S6-α-S8 core in regular all-atom simulations at a pulling speed of 1 m/s.** Time points of the secondary structure element (SSE) detachment events were collected from the pulling simulations. Histograms show the distributions of these time points. Secondary structure elements at which the unfolding begins significantly sooner in the mutant protein are colored by red ($p \leq 0.05$, Kolmogorov-Smirnov test).

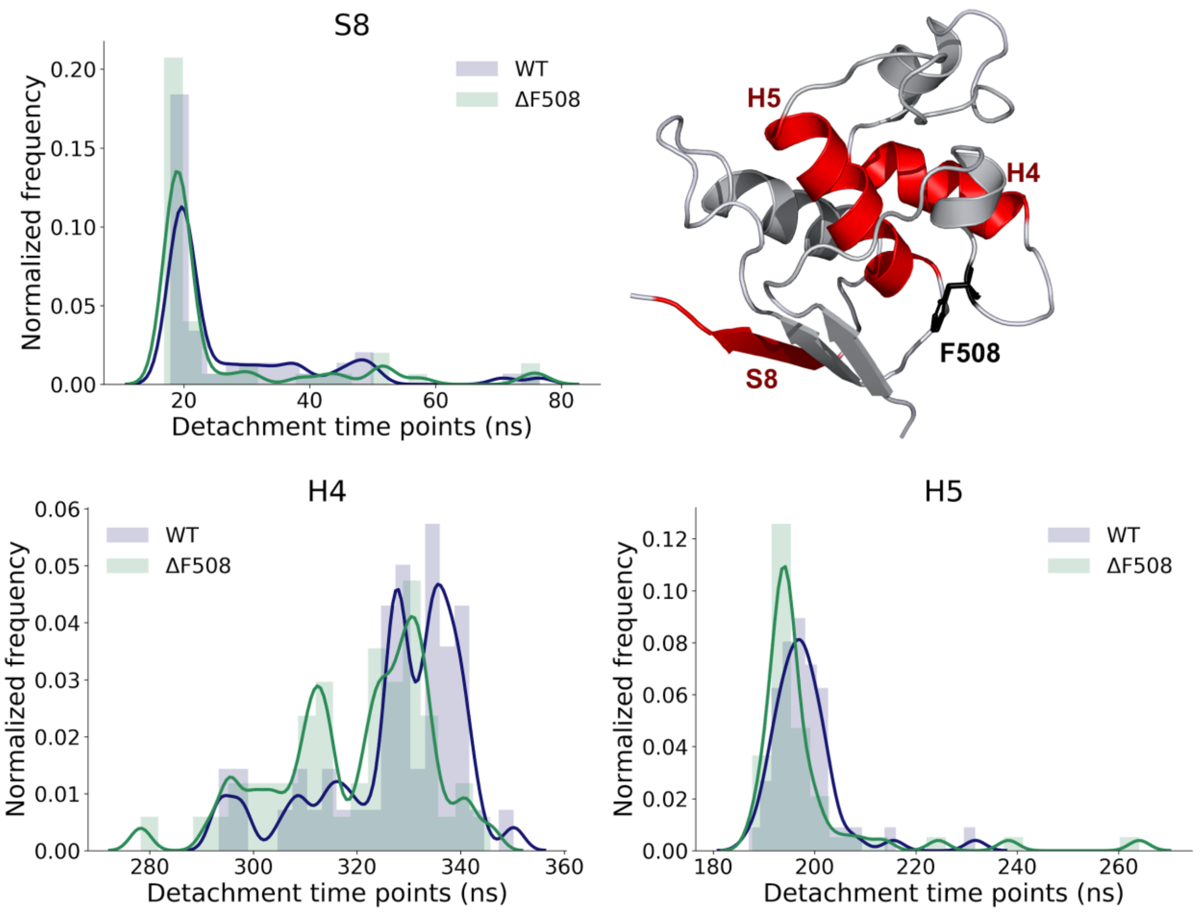


| | 1 m/s | | 0.1 m/s | |
|---|---|---|---|---|
| **SSE** | **statistic** | **p-value** | **statistic** | **p-value** |
| S6 | 0.3200 | 0.0115 | 0.1800 | 0.3959 |
| S7 | 0.1800 | 0.3959 | 0.1600 | 0.5487 |
| S8 | 0.2200 | 0.1786 | 0.3000 | 0.0217 |
| H3 | 0.2000 | 0.2719 | 0.1000 | 0.9667 |
| H4 | 0.2858 | 0.0279 | 0.3147 | 0.0102 |
| H4b | 0.0842 | 0.9861 | 0.1600 | 0.5487 |
| H5 | 0.2800 | 0.0392 | 0.3200 | 0.0115 |
| H6 | 0.2000 | 0.2719 | 0.2000 | 0.2719 |

**Supplementary Fig. 4: ΔF508 accelerates decoupling of specific secondary structure elements in the S6-α-S8 core in regular all-atom simulations also at a pulling speed of 0.1 m/s.** Time points of the secondary structure element (SSE) detachment events were collected and shown in histograms as above. Secondary structure elements at which the unfolding begins significantly sooner in the mutant protein are colored by red ($p \leq 0.05$, Kolmogorov-Smirnov test). The table shows the Kolmogorov-Smirnov test of detachment time points of selected SSE.

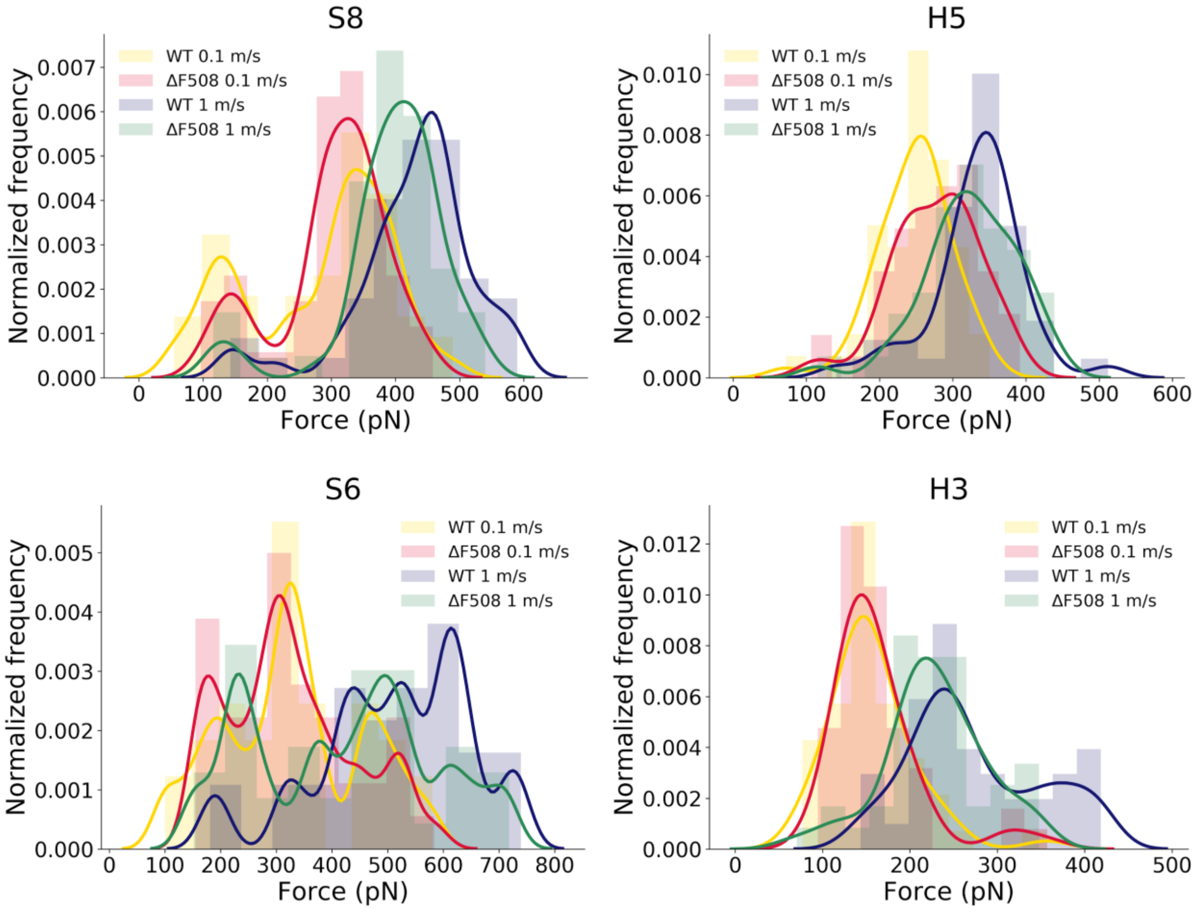


| | 1 m/s | | 0.1 m/s | |
|---|---|---|---|---|
| **SSE** | **statistic** | **p-value** | **statistic** | **p-value** |
| S6 | 0.2875 | 0.0777 | 0.1653 | 0.6074 |
| S7 | 0.3778 | 0.1485 | 0.2063 | 0.3852 |
| S8 | 0.2742 | 0.0387 | 0.1725 | 0.4023 |
| H3 | 0.2879 | 0.0581 | 0.1150 | 0.8484 |
| H4 | 0.2273 | 0.3948 | 0.1772 | 0.4462 |
| H4b | 0.2706 | 0.1559 | 0.1203 | 0.8135 |
| H5 | 0.1961 | 0.3142 | 0.3600 | 0.0028 |
| H6 | 0.1517 | 0.5617 | 0.1800 | 0.3959 |

**Supplementary Fig. 5: ΔF508 decreases the rupture forces between some secondary structural elements.** The deletion mutation accelerates the detachment by decreasing the forces between secondary structural elements at both faster and slower speeds. Some of the differences are also statistically significant (Kolmogorov-Smirnov tests are shown in the table).

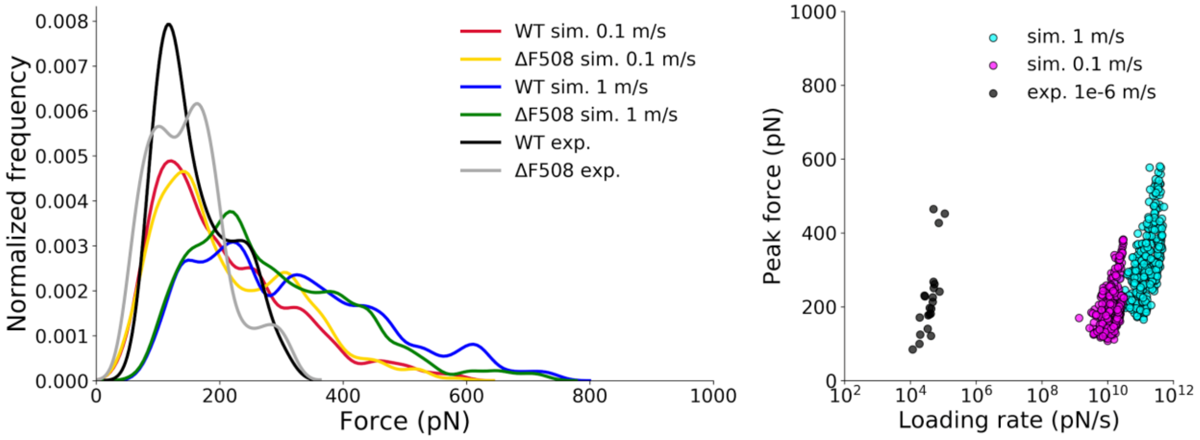


**Supplementary Fig. 6: Rupture force distribution and loading rates at various pulling speeds.** While the low pulling speed used in our experiments cannot be approached in simulations, rupture forces are in the same order of magnitude. This is likely the source of the good agreement between our experimental and computational results. The histogram shows all rupture forces belonging to the detectable events, which are not necessarily the same in simulations and experiments.

| pathway | 0.1 m/s | | 1 m/s | | 1 m/s rev. | |
|---|---|---|---|---|---|---|
| | WT | ΔF508 | WT | ΔF508 | WT | ΔF508 |
| S6 S7-S8 H6 H3-H5 H4 | | | | | | 2% |
| S6 S7-S8 H6 H5 H3 H4 | 10% | 8% | 4% | | 2% | |
| S6 S8 H6-S7 H5 H3 H4 | | | | | 2% | |
| S6 S8 S7 H6 H3 H5 H4 | | | 2% | | | 2% |
| S6 S8 S7 H6 H5 H3 H4 | 12% | 10% | 4% | 6% | | 10% |
| S6 S8 S7 H6 H5 H4 H3 | | 2% | | | | |
| S6-S7-S8 H6 H5 H3 H4 | 4% | | | | | |
| S6-S8 S7 H6 H3-H5 H4 | | | | | | 2% |
| S6-S8 S7 H6 H5 H3 H4 | 4% | | | 4% | | 2% |
| S7-S8 S6 H6 H5 H3 H4 | 2% | | | | | |
| S8 H6 S6 S7 H5 H3 H4 | | 2% | | 6% | | 2% |
| S8 H6 S6-S7 H3 H5 H4 | | 4% | 4% | 4% | 8% | 4% |
| S8 H6 S6-S7 H3-H5 H4 | 4% | 2% | 20% | 6% | 2% | 8% |
| S8 H6 S6-S7 H5 H3 H4 | 44% | 52% | 54% | 44% | 76% | 48% |
| S8 H6 S6-S7 H5 H3-H4 | | 2% | | | 2% | 4% |
| S8 H6 S6-S7 H5 H4 H3 | 2% | | | | | |
| S8 H6-S6-S7 H5 H3 H4 | 4% | | | | | |
| S8 S6 S7 H5-H6 H3 H4 | | | | 2% | | |
| S8 S6 S7 H6 H3 H5 H4 | | | 2% | 2% | | |
| S8 S6 S7 H6 H5 H3 H4 | | 2% | | 8% | | 4% |
| S8 S6-S7 H6 H3 H5 H4 | 2% | 2% | | 4% | | 2% |
| S8 S6-S7 H6 H5 H3 H4 | 10% | 14% | 10% | 14% | 6% | 10% |
| S8 S7 S6 H6 H5 H3 H4 | 2% | | | | 2% | |

**Supplementary Fig. 7: Unfolding pathways of the S6-α-S8 core region in the case of pulling velocities of 0.1 m/s, 1 m/s and in reversed direction (1 m/s rev).** Pathways were determined by the sequence of detachment events of secondary structure units. Synchronized unfolding of two elements is marked by hyphenation. Most frequent pathway was marked with a blue background.

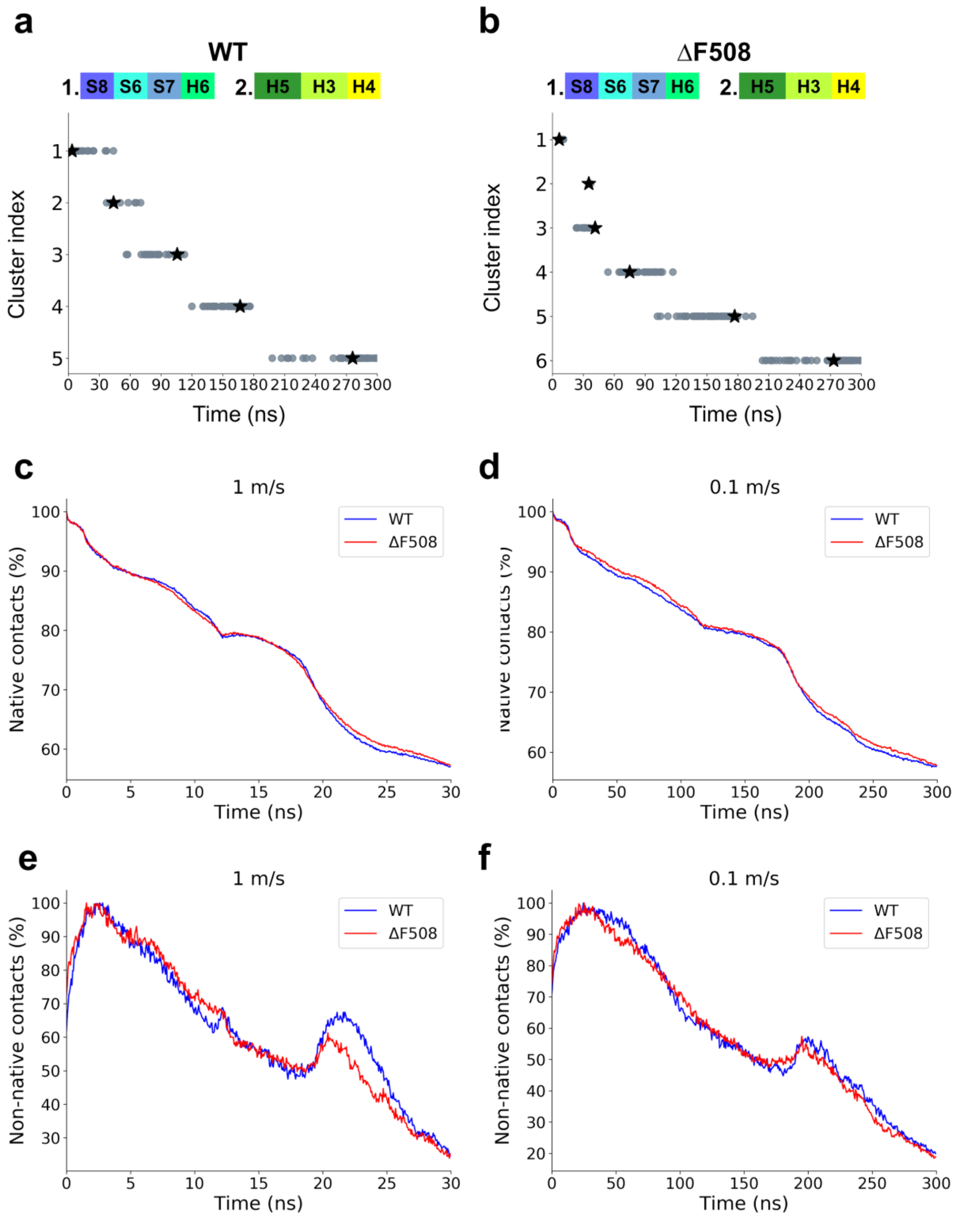


**Supplementary Fig. 8: Clusters and contact changes of S6-α-S8 core unfolding.** (**a, b**) Intermediate structures from all WT and ΔF508 pulling simulations at velocity 0.1 m/s were clustered using contact RMSD. Cluster centroids are indicated by stars. The number of native **(c, d)** and non-native **(e, f)** contacts were calculated, normalized to their maximum value, and plotted at pulling velocities indicated in the plots.

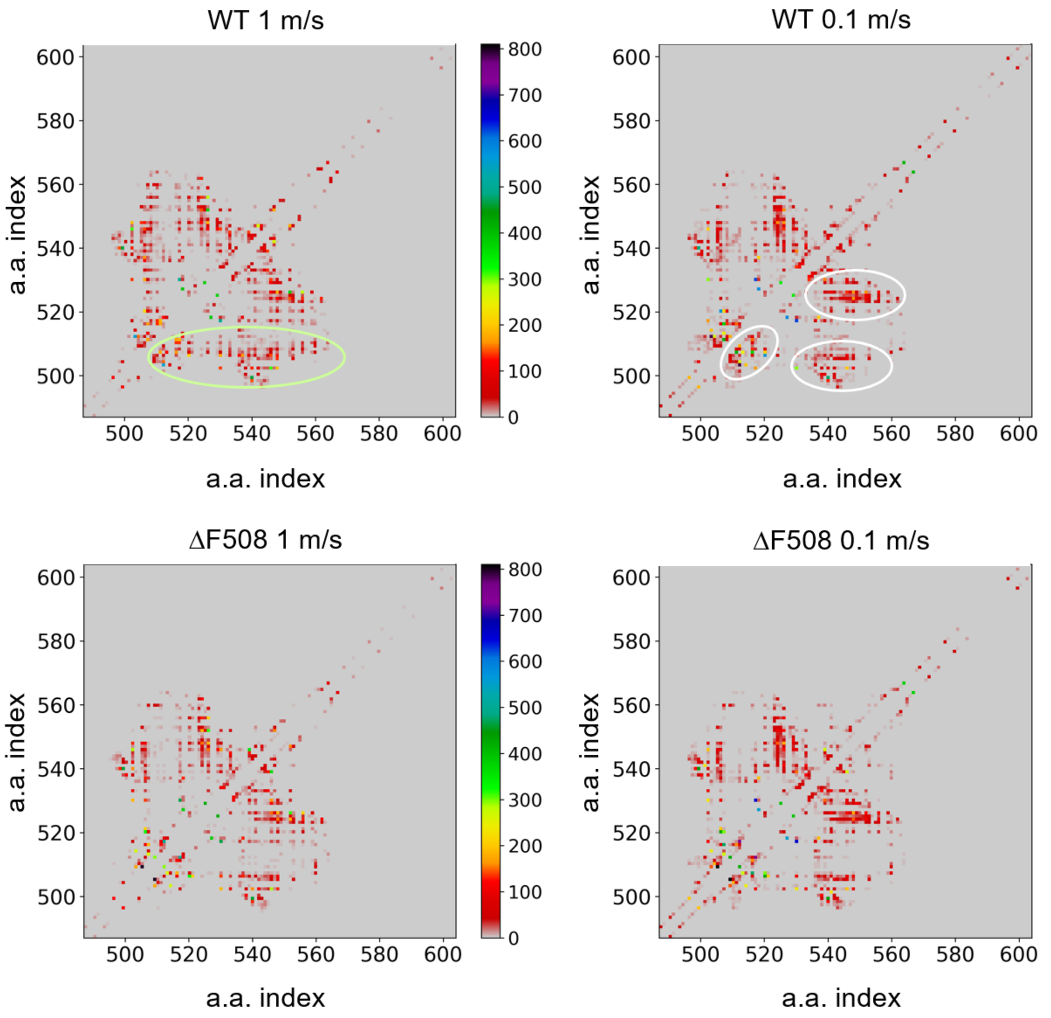


**Supplementary Fig. 9: Proximity values were used to identify important non-native contacts in unfolding simulations.** Summed proximity values, which were calculated for all the WT and the ΔF508 simulations (n=50 for each panel) carried out with pulling velocities of 1 m/s and 0.1 m/s, were plotted. Proximity matrices show the non-native contacts between 18-25 ns and 180-250 ns at velocities of 1 m/s and 0.1 m/s, respectively. The green border indicates the scattered interacting residues of F508 at 1 m/s. White borders indicate the more localized interacting residues of F508 at 0.1 m/s.

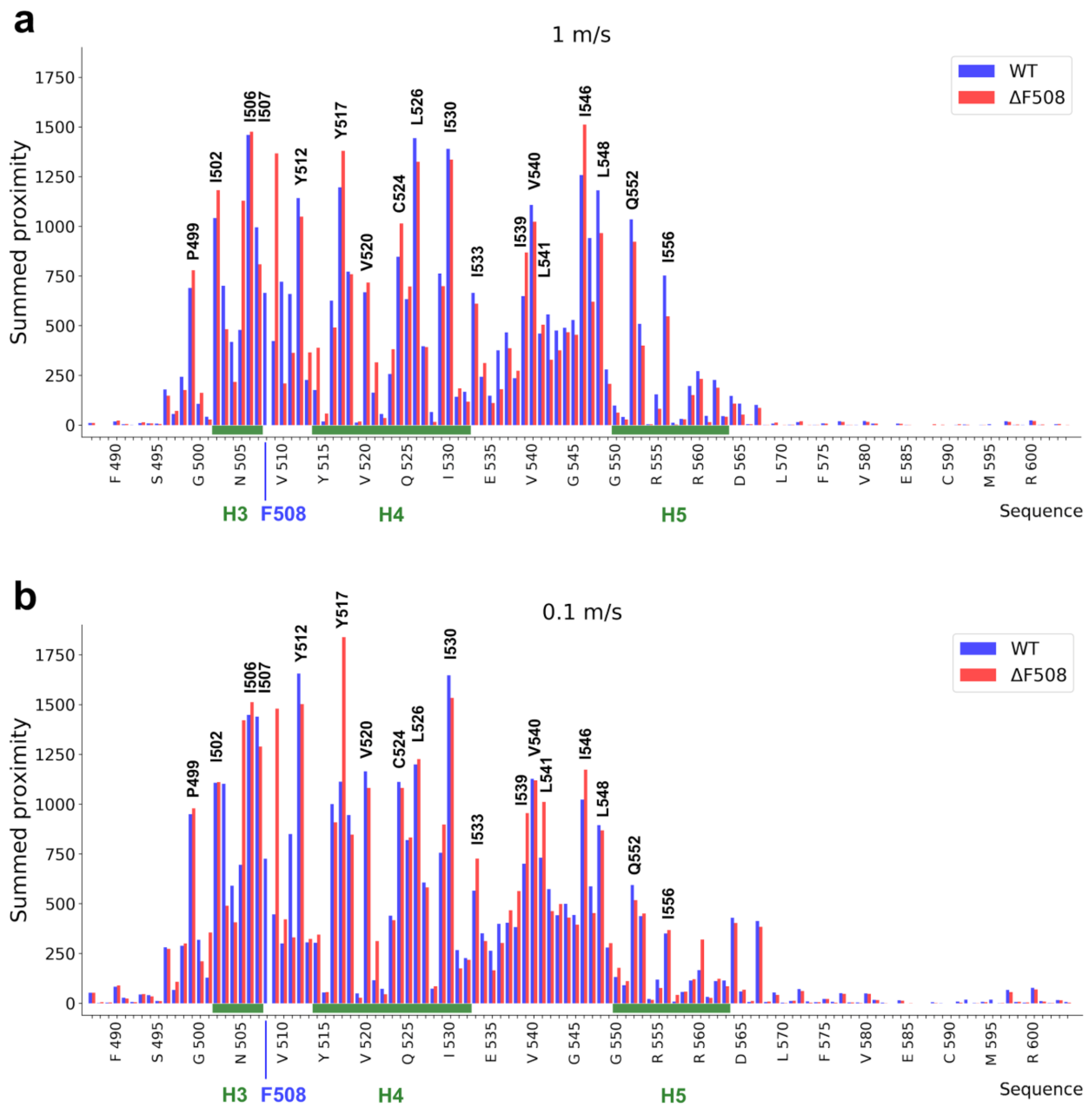


**Supplementary Fig. 10: Summed 2D proximity values in pulling simulations.** Proximity values of non-native contacts during the trajectory along the sequence of S6-α-S8 core were calculated and summed for each residue in simulations performed with pulling velocity of 1 m/s (**a**) and 0.1 m/s (**b**). Higher values indicate that the residue has many close interactions during the investigated time period of unfolding (18-25 ns) which interactions are not present in the native structure. Blue and red columns represent the wild type and the ΔF508 mutant, respectively.

**a**

**WT**

| a. a. # | | proximity |
|---|---|---|
| I | 506 | 1460 |
| L | 526 | 1443 |
| I | 530 | 1389 |
| I | 546 | 1257 |
| Y | 517 | 1196 |
| L | 548 | 1180 |
| Y | 512 | 1142 |
| V | 540 | 1107 |
| I | 502 | 1041 |
| Q | 552 | 1035 |
| I | 507 | 995 |
| T | 547 | 941 |
| C | 524 | 846 |
| R | 518 | 772 |
| D | 529 | 762 |
| I | 556 | 753 |
| V | 510 | 721 |
| K | 503 | 701 |
| P | 499 | 689 |
| V | 520 | 668 |
| F | 508 | 665 |
| F | 533 | 665 |
| S | 511 | 659 |
| I | 539 | 649 |
| Q | 525 | 633 |
| R | 516 | 626 |
| G | 542 | 556 |
| G | 545 | 528 |
| R | 553 | 509 |

**b**

**ΔF508**

| a. a. # | | proximity |
|---|---|---|
| I | 546 | 1511 |
| I | 506 | 1476 |
| Y | 517 | 1379 |
| G | 509 | 1367 |
| I | 530 | 1335 |
| L | 526 | 1325 |
| I | 502 | 1182 |
| N | 505 | 1128 |
| Y | 512 | 1049 |
| V | 540 | 1023 |
| C | 524 | 1014 |
| L | 548 | 965 |
| Q | 552 | 922 |
| I | 539 | 868 |
| I | 507 | 808 |
| P | 499 | 778 |
| R | 518 | 759 |
| V | 520 | 717 |
| D | 529 | 698 |
| Q | 525 | 697 |
| T | 547 | 621 |
| F | 533 | 611 |
| I | 556 | 547 |
| L | 541 | 504 |

**c**

**ΔF508-WT**

| a. a. # | | Proximity difference |
|---|---|---|
| G | 509 | 945 |
| F | 508 | -665 |
| N | 505 | 649 |
| V | 510 | -512 |
| T | 547 | -320 |
| S | 511 | -297 |
| I | 546 | 254 |
| G | 542 | -228 |
| I | 539 | 220 |
| K | 503 | -219 |
| L | 548 | -215 |
| E | 514 | 213 |
| I | 556 | -206 |
| E | 504 | -201 |
| K | 536 | -195 |
| I | 507 | -186 |
| Y | 517 | 183 |
| C | 524 | 167 |
| I | 521 | 153 |
| I | 502 | 141 |
| D | 513 | 138 |
| R | 516 | -135 |
| A | 523 | 124 |
| L | 526 | -118 |
| Q | 552 | -113 |
| R | 553 | -109 |
| E | 543 | -100 |

**Supplementary Fig. 11:** (**a**, **b**) Amino acid residues with high proximity values (>500) for both wild type and ΔF508 mutant S6-α-S8 core from pulling simulations (1 m/s) and (**c**) amino acid residues where the absolute value of the proximity difference of the mutant and wild type is above a threshold: abs(proximity difference) > 100. Colored fields indicate residues participating in the non-native contact formation that are characteristic either of the wild type (blue) or the ΔF508 mutant (red).

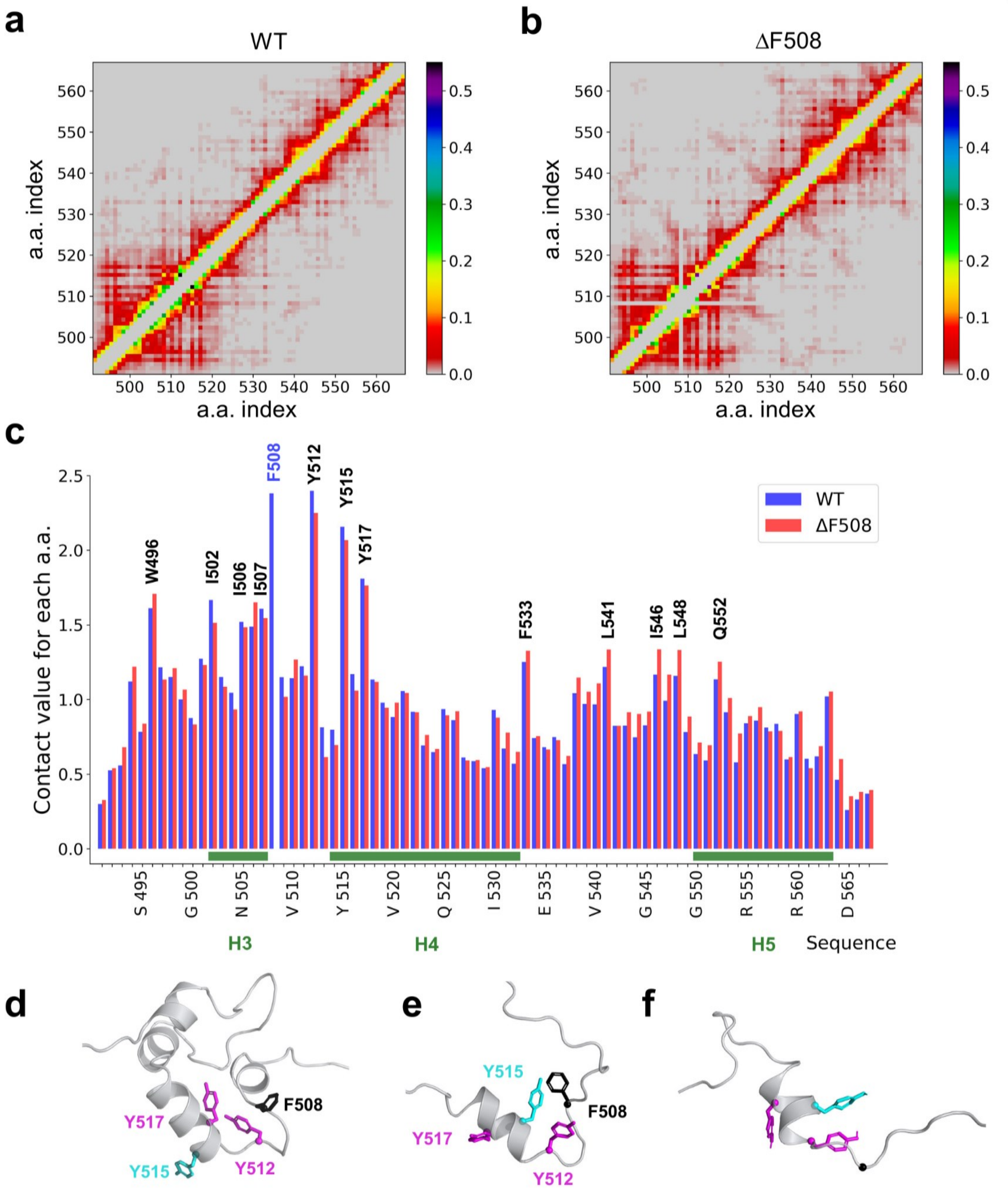


**Supplementary Fig. 12: NBD1 folding REX-DMD simulations.** (**a**) Contact frequency of pairwise amino acid contacts observed in conformations, which visited the highest temperature in replica exchange DMD folding simulations, were calculated and plotted. (**b**) Summed contact values of each amino acid were calculated and plotted. (**d**) Conformation of hydrophobic amino acids in the natively folded NBD1. (**e**) F508-Y515 is a frequent non-native interaction in folding simulations and may provide an important intermediate step during NBD1 folding. (**f**) The lack of F508 results in a lower probability of turn formation by the a.a. 508-512 region that likely affects NBD1 folding adversely.

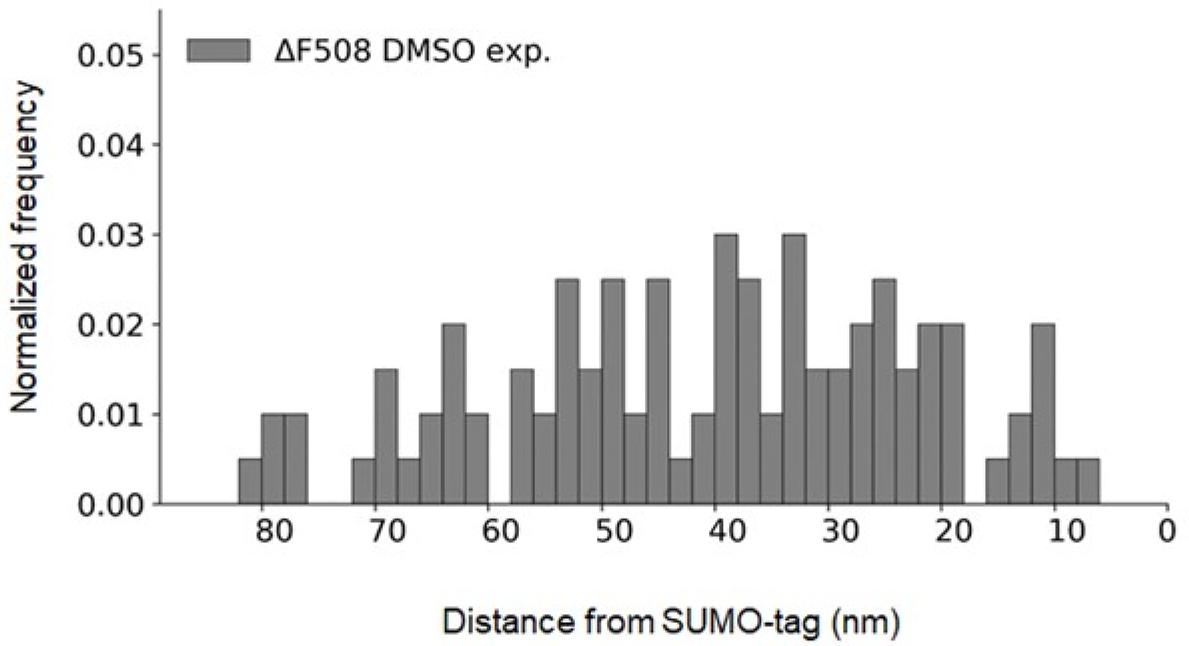


**Supplementary Fig. 13: The distribution of contour lengths increases from ΔF508 mutant in the presence of DMSO.** The solvent of BIA did not have an effect of unfolding.

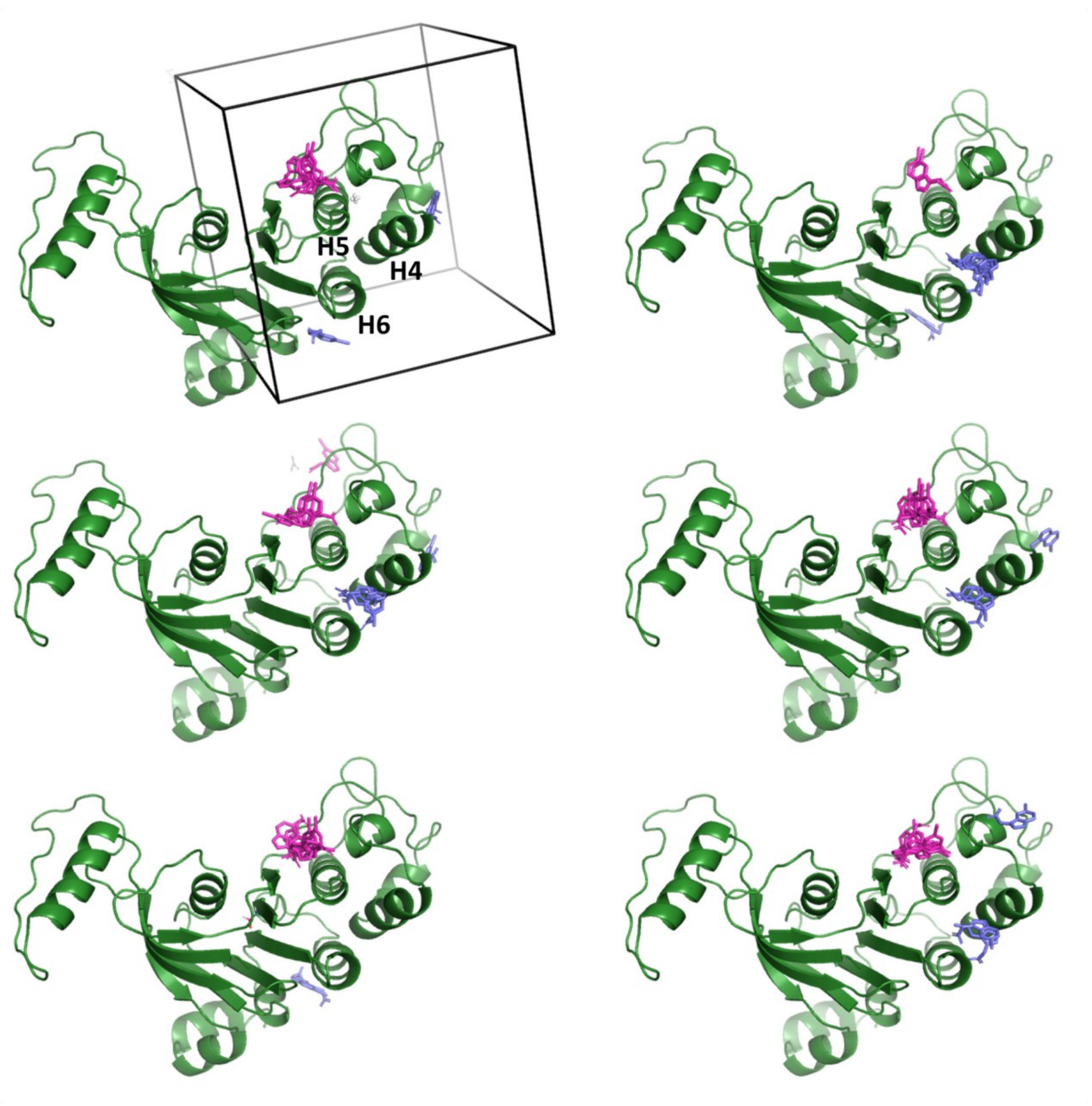


**Supplementary Fig. 14: *In silico* docking suggests H4, H5, and H6 as the BIA binding site.** BIA was docked to six equilibrated NBD1 structures from MD simulations using Autodock Vina. When the full NBD1 was included in the search box, a major docking site was the regulatory insertion. Because its dynamics was not altered in the presence of BIA (Supplementary Fig. 15), we included only the α-subdomain into the search space (black box). In this case, one of the major binding sites was the CL4 pocket, which include BIA decoys (magenta sticks), while the second most populated site involves H4, H5, and H6 helices, intercalating BIA molecules (blue sticks).

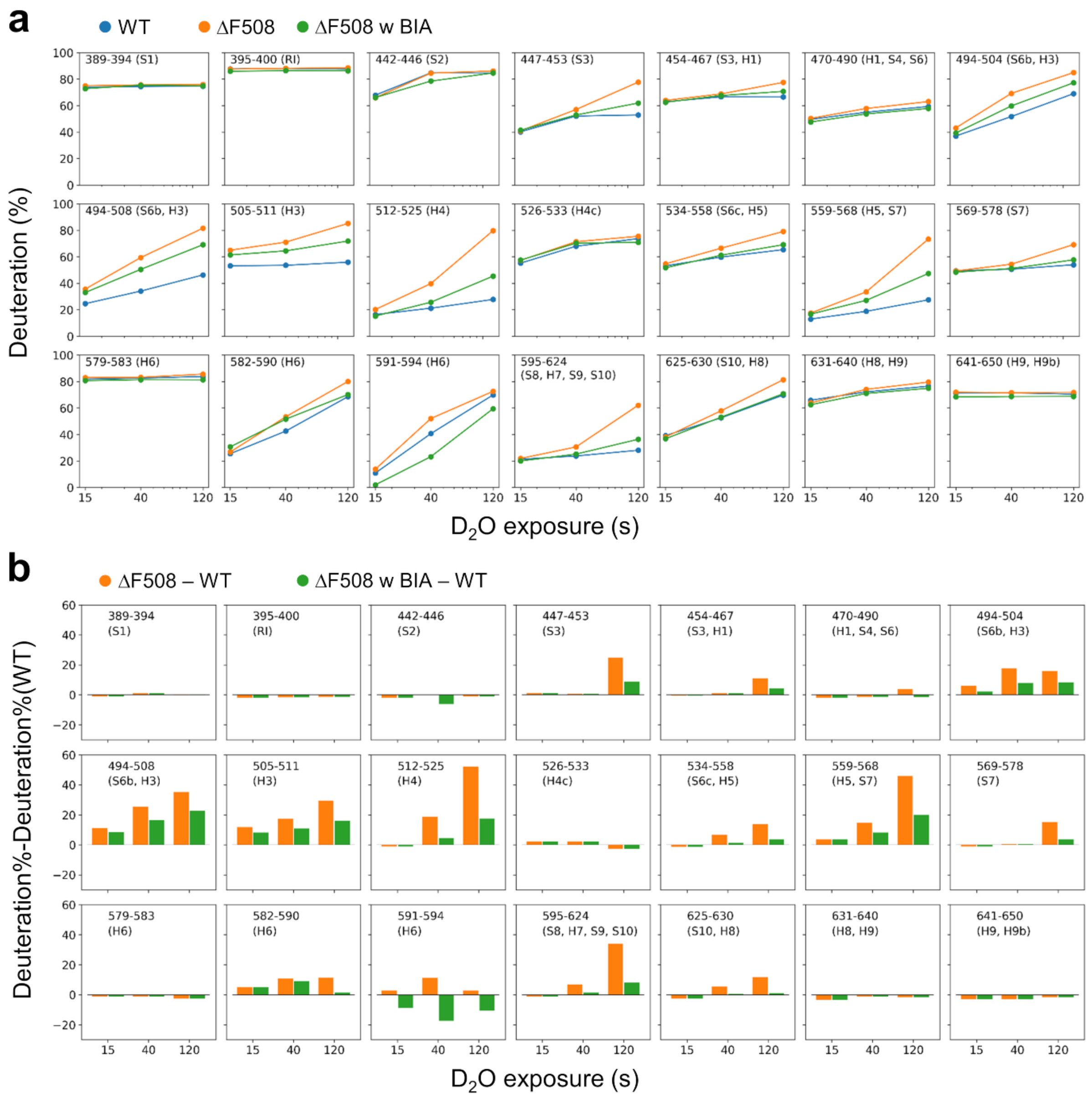


**Supplementary Fig. 15: HDX kinetics of peptides corresponding to isolated WT- and ΔF508-NBD1 in the absence and presence of BIA.** (**a**) HDX% were measured for 15, 40, and 120 s (n>=3) and plotted. (**b**) In order to highlight the changes upon BIA addition, we also plotted the deuteriation levels compared to the WT level. Some ΔF508-NBD1 regions (447-453, 512-525, 559-568, 591-594, and 595-624) exhibited decreased dynamics upon BIA binding. The segment 591-594 had decreased dynamics even at the shortest time point, strongly suggesting that the backbone NH exchange was directly protected by BIA rather than via allosteric effects as observed for distant peptides of the ΔF508-NBD1.

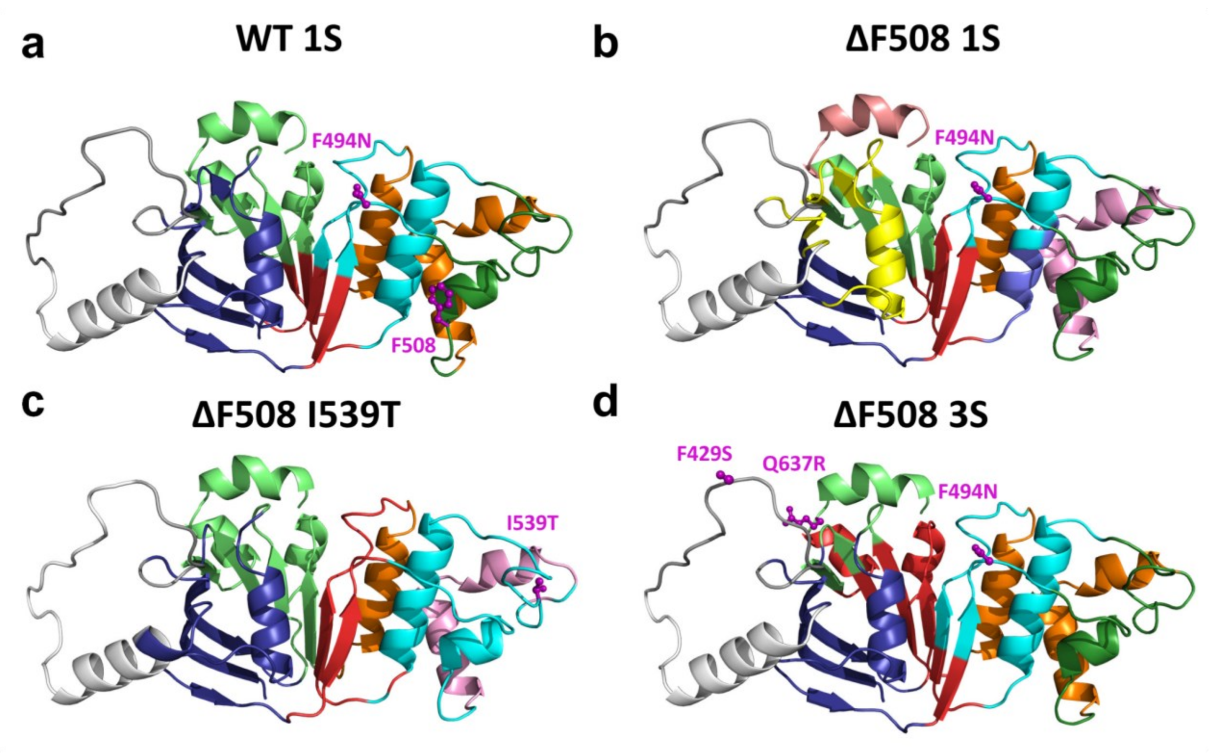


**Supplementary Fig. 16: Symmetricity of allosteric coupling within NBD1.** (**a, b**) ΔF508 disrupted dynamic communities (groups of residues with similar motions, labeled by different colors) both in the α- and β-subdomains, observed in the WT. E.g. dynamics of the H1 helix in ΔF508 (yellow) was decoupled from S1, S2, and S4 strands (blue). (**c**) The I539T secondary site mutation, which is located in the α-subdomain, highly increased the dynamic coupling of the α-subdomain. This is indicated by restoration of H5 helix as a single community (cyan) and inclusion of H3 (green in WT) into this community. Importantly, I539T also restored the dynamic communities in the β-subdomain. (**d**) Two additional secondary site mutations (F429S and Q637R), which are located in the β-subdomain, rescued not only the community pattern observed in the WT 1S β-subdomain, but also in the α-subdomain. Moreover, they increased the coupling between S3, S5, S6, S7, and S8, forming a single community (red). H4a, H4b, and H6 helices were also merged into one dynamic community (orange). These results support a strong and symmetric allosteric communication between subdomains.

| ∆L | WT | ∆F508 |
|---|---|---|
| 12nm → 30nm | 31.3% | 14.7% |
| 30nm → 12nm | 16.7% | 8.8% |
| 42nm | 8.3% | 4.4% |
| | **56.3%** | **27.9%** |

**Table S1: The frequency of S6-α-S8 core unfolding patterns from AFM pulling experiments.**
